# Celda: A Bayesian model to perform co-clustering of genes into modules and cells into subpopulations using single-cell RNA-seq data

**DOI:** 10.1101/2020.11.16.373274

**Authors:** Zhe Wang, Shiyi Yang, Yusuke Koga, Sean E. Corbett, W. Evan Johnson, Masanao Yajima, Joshua D. Campbell

## Abstract

Single-cell RNA-seq (scRNA-seq) has emerged as a powerful technique to quantify gene expression in individual cells and elucidate the molecular and cellular building blocks of complex tissues. We developed a novel Bayesian hierarchical model called Cellular Latent Dirichlet Allocation (Celda) to perform simultaneous co-clustering of genes into transcriptional modules and cells into subpopulations. Celda can quantify the probabilistic contribution of each gene to each module, each module to each cell population, and each cell population to each sample. We used Celda to identify transcriptional modules and cell subpopulations in a publicly available peripheral blood mononuclear cell (PBMC) dataset. Celda identified a population of proliferating T cells and a single plasma cell which were missed by two other clustering methods. Celda identified transcriptional modules that highlighted unique and shared biological programs across cell types. Celda also outperformed a PCA-based approach for gene clustering on simulated data. Overall, Celda presents a novel statistically principled approach towards characterizing transcriptional programs and cellular heterogeneity in single-cell RNA-seq data.

## Introduction

Complex biological systems can be conceptually defined into hierarchies where each level of the hierarchy is composed of different subunits which cooperate to perform distinct biological functions^1^. For example, organisms can be subdivided into a collection of complex tissues: each complex tissue is composed of different cell types; each cell population is denoted by a unique combination of transcriptionally activated pathways (i.e. transcriptional modules); and each transcriptional module is composed of genes that are coordinately expressed to perform specific molecular functions. By identifying the “building blocks” and their composition within each level of the hierarchy, we can more readily identify the patterns that define the behavior of these elements.

Single-cell RNA-seq (scRNA-seq) is a molecular assay that can quantify gene expression patterns in individual cells. In contrast to profiling of “bulk” RNA from a sample, where only an average transcriptional signature across all the composite cells can be derived, scRNA-seq experiments can profile thousands of single-cell transcriptomes per sample and thus offer an excellent opportunity to identify novel subpopulations of cells and to characterize transcriptional programs that define each subpopulation by examining co-varying patterns of gene expression across cells^2^. However, analysis of scRNA-seq data presents several challenges. For example, the data tends to be sparse due to the difficulty in amplifying low amounts of RNA in individual cells. To combat noise from the amplification process, unique molecular identifiers (UMIs) are often incorporated to eliminate duplicate reads derived from the same mRNA molecule^3^. The use of these UMIs enables the measurement of discrete counts of mRNA transcripts within each cell, making models constructed using discrete distributions a suitable approach for analyzing this type of data.

Discrete Bayesian hierarchical models have proven to be powerful tools for unsupervised modeling of discrete data types. In the text mining field, a plethora of models have been developed that can identify hidden topics across documents and/or cluster documents into distinct groups^4-8^. These models generally treat each document as a “bag-of-words” where each document is represented by a vector of counts or frequencies for each word in the vocabulary. Each document cluster (hidden topic) is represented by a Dirichlet distribution where words with higher probability are observed more frequently for the document cluster^5^. Given the success of topic models with sparse text data, and the discrete, sparse nature of transcriptional data generated by many scRNA-seq protocols, the application of such discrete Bayesian hierarchical models represents a promising approach to characterize structures in scRNA-seq data.

Various scRNA-seq clustering workflows and methods including ascend^9^, BAMM-SC^10^, CIDR^11^, DESC^12^, DIMM-SC^13^, pcaReduce^14^, SAFE-clustering^15^, SAME-clustering^16^, SC3^17^, scran^18^, Seurat^19^, SIMLR^20^, TSCAN^21^, and VPAC^22^ have been developed. However, methods that can group genes into transcriptional modules to elucidate the combination of underlying biological programs that define each cell cluster have not been reported. We developed a model (**Celda_CG**) that simultaneously perform co-clustering of <u>**C**</u>ells into subpopulations and <u>**G**</u>enes into transcriptional modules. While this model can perform clustering of genes and/or cells, it also has the ability to describe the relationship between different layers of a biological hierarchy via probabilistic distributions. These distributions constitute dimensionally reduced representations of the data that can be used for down-stream exploratory analysis. We demonstrate the utility of this approach by applying the Celda_CG model to a publicly available scRNA-seq dataset of peripheral blood mononuclear cells (PBMCs). Celda_CG identifies novel cell subpopulations missed by other approaches while characterizing transcriptional programs that are active to various degrees within and across major cell types.

## Results

We developed a novel discrete Bayesian hierarchical model, called <u>Ce</u>llular <u>L</u>atent <u>D</u>irichlet <u>A</u>llocation (Celda), to simultaneously perform co-clustering of genes into modules and cells into subpopulations (**Figure 1, Supplementary Information**). Each level in the biological hierarchy is modeled as a mixture of components using Dirichlet distributions: sample *i* is a mixture of cellular subpopulations (*θ*_*i*_), each cell subpopulation *k* is a mixture (*φ*_*k*_) of transcriptional modules, and each module *l* is a mixture (*ψ*_*l*_) of features such as genes. *θ*_*i,k*_ is the probability of cell population in *k* sample *i, φ*_*k,l*_ is the probability of module *l* in population *k*, and *ψ*_*l,g*_ is the probability of gene *g* in module *l* (**Figure 1a, b**). Each cell *j* in sample *i* has a hidden cluster label, *z*_*i,j*_ denoting the population to which it belongs. Each transcript *x*_*i,j,t*_, has a hidden label *w*_*i,j,t*_ denoting the transcriptional module to which it belongs. A similarly structured topic model has previously been proposed called “Latent Dirichlet Co-Clustering”8. However, we add a unique and novel component to our model specifically geared towards gene expression analysis.

**Figure 1.**
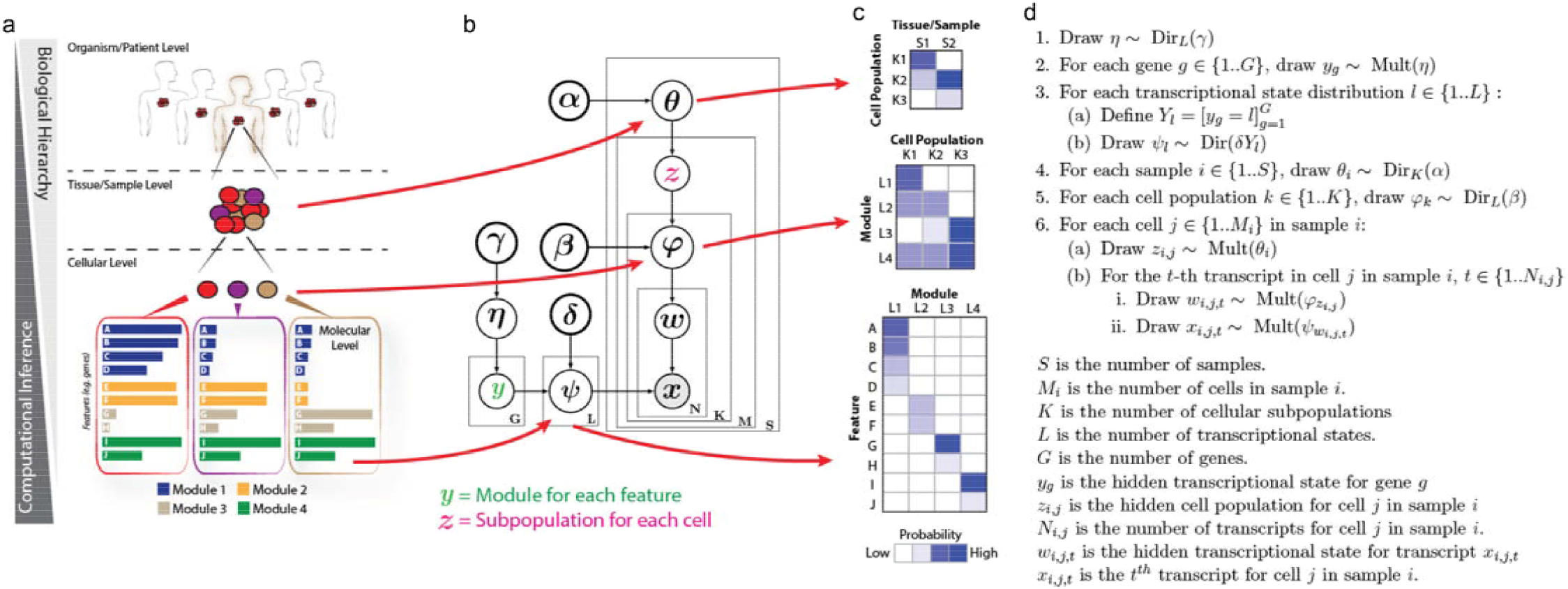
Celda identifies cell heterogeneity by clustering genes into modules and cells into subpopulations. **a**, Example of a biological hierarchy. One way in which we try to understand complex biological systems is by organizing them into hierarchies. Individual organisms are composed of complex tissues. Each complex tissue is composed of different cellular populations with distinct functions; each cellular subpopulation contains a unique mixture of molecular pathways (i.e. modules); and each module is composed of groups of genes that are co-expressed across cells. **b**, Plate diagram of Celda_CG model. We developed a novel discrete Bayesian hierarchical model called Celda_CG to characterize the molecular and cellular hierarchies in biological systems. Celda_CG performs “co-clustering” by assigning each gene to a module and each cell to a subpopulation. **c**, In addition to clustering, Celda_CG also inherently performs a form of “matrix factorization” by deriving three distinct probability matrices: 1) a Cell Population x Sample matrix representing the probability that each population is present in each sample, 2) a Transcriptional Module x Cell Population matrix representing the contribution of each transcriptional state to each cellular subpopulation, and 3) a Gene x Module matrix representing the contribution of each gene to its Module. **d**, Generative process for the Celda_CG model.

The goal of many gene expression clustering algorithms is to group genes into distinct, non-overlapping sets of genes^23-26^ (i.e. hard-clustering of genes). The rationale for this type of clustering is that genes that co-vary across cells and samples are likely involved in the same biological processes and should be considered a single biological program^27^. In order to enforce “hard-clustering” of genes into modules, we modified an approach from Wang and Blei^6^ regarding the sparse Topic Model (sparseTM) that has the capability to turn words “on” or “off” in different topics, by assigning a non-zero or zero probability to that word in each topic. In Celda_CG, we leverage this technique to turn off genes in all modules except one to enable the hard-clustering behavior.

While Celda can perform clustering, it also offers probabilistic distributions which describes the contribution of each “building block” to each layer of the biological hierarchy (**Figure 1c**). These distributions can also be viewed as reduced dimensional representations of the data that can be used for downstream exploratory analyses. For example, the *φ* matrix contains the probability of each module in each cell population and thus provides a high-level view of the structure of the dataset. In addition to Celda_CG, we have also developed two distinct models that cluster <u>**C**</u>ells into subpopulations (**Celda_C**) or cluster <u>**G**</u>enes into transcriptional modules (**Celda_G, Supplementary Methods**).

### Identification of cell populations in PBMCs

To assess Celda_CG’s ability to identify biologically meaningful cell subpopulations in real-world scRNA-seq data, we applied it to a publicly available dataset provided by 10X Genomics. The dataset (PBMC 4K) contains 4,340 PBMCs collected from a healthy donor. To determine the optimal number of transcriptional modules (*L*) and cell populations (*K*), we employed a step-wise splitting procedure first for the number of modules using a temporary cell-clustering solution and then for the number of cell populations using a fixed number of modules (**Methods, Supplementary Figure 1**). The rate of perplexity change^28^ (RPC) was measured at each split. An RPC closer to zero indicates that the addition of new modules or cell clusters is not substantially decreasing the perplexity. By observing the “elbows” on these curves as a reference point in combination with manual review of module heatmaps and cell clusters, a solution of *L =* 80 transcriptional modules and *K =* 20 cell populations the “elbows” on these curves as a reference point in combination with manual review of module was chosen for further characterization.

A UMAP^29^ dimension reduction representation was generated based on the estimated module probabilities for each cell and the major subtypes of immune cells were identified by examining expression of known marker genes (**Figure 2, Supplementary Table 1**). Among the 20 identified cell clusters in the PBMC sample, we identified major immune cell populations including CD19^+^ B-cells, FCER1A^+^ dendritic cells (DCs), CLEC4C^+^ plasmacytoid dendritic cells (pDCs), CD34^+^ progenitor cells, KLRD1^+^ natural killer cells (NKs), ITGA2B^+^ megakaryocytes, CD14^+^ monocytes, FCGR3A^+^ monocytes, CCR7^+^ memory T-cells, CD8A^+^CD8B^+^ cytotoxic T-cells, and CD4^+^ T helper cells. Cell subpopulations 15–20 show a consistently higher expression (FDRs < 0.01) of the T cell marker genes CD3D, CD3E, and CD3G relative to all other clusters. Among these T cell subpopulations. Clusters 17, 18, and 19 show consistently higher expressions (FDRs < 0.01) of CD8A and CD8B (**Supplementary Figure 2**). Within these CD8A^+^CD8B^+^ T cells, cluster 17 has high expression (FDR < 0.01) of naive T cell marker CCR7, whereas cluster 18 has consistent high expressions (FDRs < 0.01) of NK cell markers GNLY, KLRG1, and granzyme genes GZMA and GZMH, so we classified them as naive CD8^+^ T-cells and NKT cells, respectively^30^. Cell subpopulation 15 expressed T-cell markers as well as uniquely high levels of module 61, which contained genes associated with proliferation including MKI67, IL2RA, CENPM, and CENPF (**Supplementary Figure 2**) commonly found in activated proliferating T-cells^31, 32^. Cell subpopulation 1 contained a single cell which had the highest number of UMIs across the dataset (n = 48,443). This cell expressed several B lineage markers such as CD79A, CD79B, and CD19 but also contained a relatively high fraction (27%) of UMIs for immunoglobulin heavy chain and light chain genes IGHG1, IGHG3, IGLC2 and IGLC3. These genes were not observed in other cells and suggest a plasma cell lineage^33^ (**Supplementary Figure 3**). Cell populations 1 and 15 were not identified by the analysis workflows and graph-based clustering methods used in Seurat^19^ and scran^18^ packages (**Supplementary Figure 4**), demonstrating Celda’s ability to characterize additional cellular heterogeneity compared to other methods.

**Figure 2.**
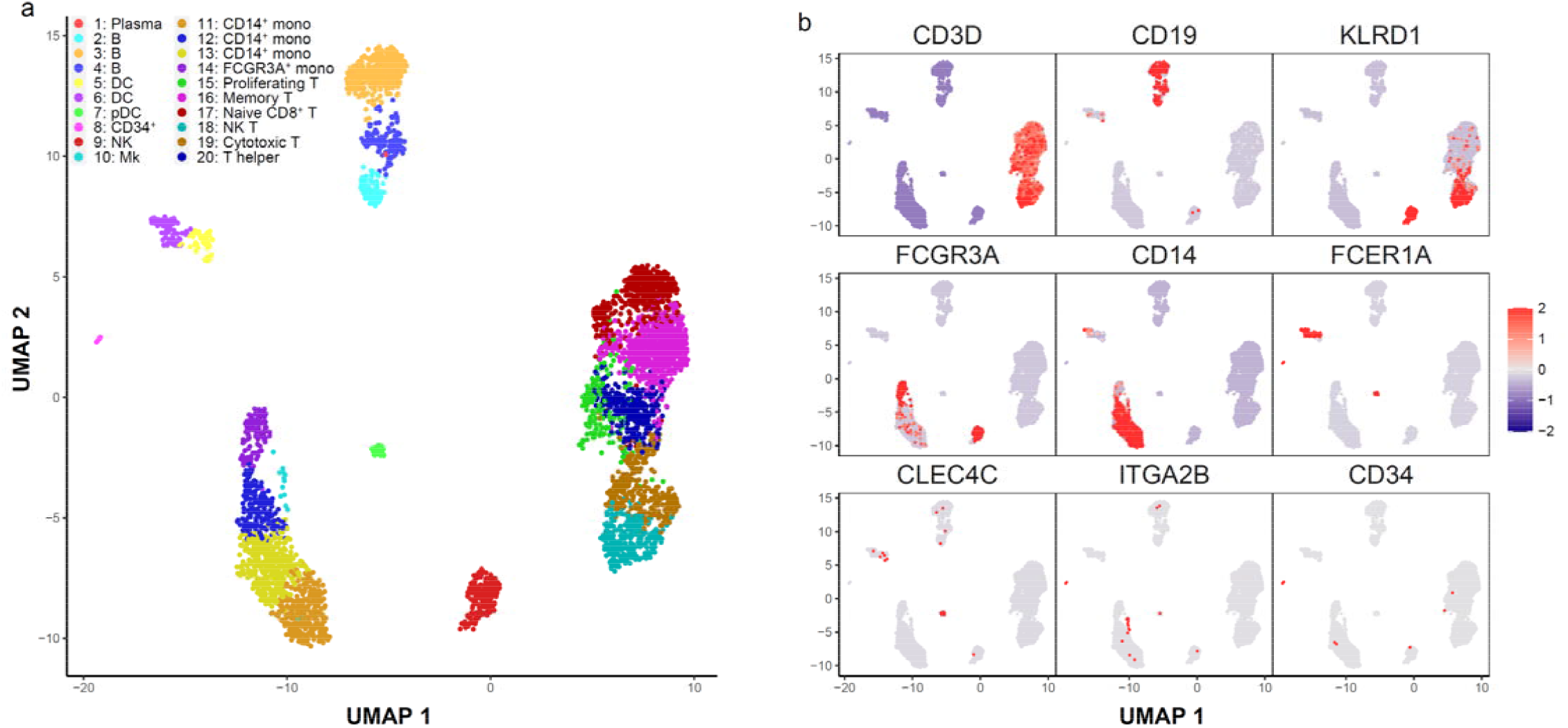
Celda identifies immune cell subpopulations from PBMC scRNA-seq data. To demonstrate the utility of Celda clustering model, we applied it to a scRNA-seq dataset of 4,340 peripheral blood mononuclear cells (PBMCs) generated using 10X Chromium platform and identified 80 transcriptional modules and 20 cell populations. **a**, UMAP dimension reduction representation of 4,340 PBMCs based on the transcriptional module probabilities. **b**, Scaled normalized expressions of representative gene markers show clustering of cell subpopulations including T cells (CD3D), B cells (CD19), natural killer cells (KLRD1), FCGR3A+ monocytes (FCGR3A), CD14+ monocytes (CD14), dendritic cells (FCER1A), plasmacytoid dendritic cells (CLEC4C), megakaryocytes (ITGA2B), and CD34+ progenitor cells (CD34). Cell populations 1 (proliferating T-cells) and 15 (plasma cell) are novel cell clusters identified by Celda, demonstrating Celda’s ability to characterize additional cellular heterogeneity.

### Identification of transcriptional modules with unique patterns of expression across cell populations

Beyond assessment of individual marker genes, Celda has the ability to identify modules of co-expressed genes which can be further examined to characterize transcriptional programs active in one or more cell populations (**Figure 3**). An overview of the relationships between modules and cell subpopulations can be explored with the *φ* probability matrix which contains the probability of each module within each cell subpopulation (**Figure 3a**). This matrix gives insights into the absolute abundance of each module within the same cell subpopulation. For example, module 62 contains actin-related housekeeping genes such as ACTB and ARPC1B and has higher expression than most other modules within each cell population. A relative probability heatmap can also be produced by taking the z-score of the module probabilities across cell subpopulations (**Figure 3b**). Examining the relative abundance of a transcriptional module among different cell populations can be useful for finding modules that exhibit specific patterns across cell populations even if they have an overall lower absolute probability compared to other modules. For example, module 65 contains CD8A and CD8B and has an overall lower abundance compared to other housekeeping modules such as module 62 within each cell population (as can be observed in **Figure 3a**). However, module 65 has higher relative expression in the T-cell populations 17, 18, and 19 and can be used to classify CD8^+^ subpopulations (as can be observed in **Figure 3b)**.

**Figure 3.**
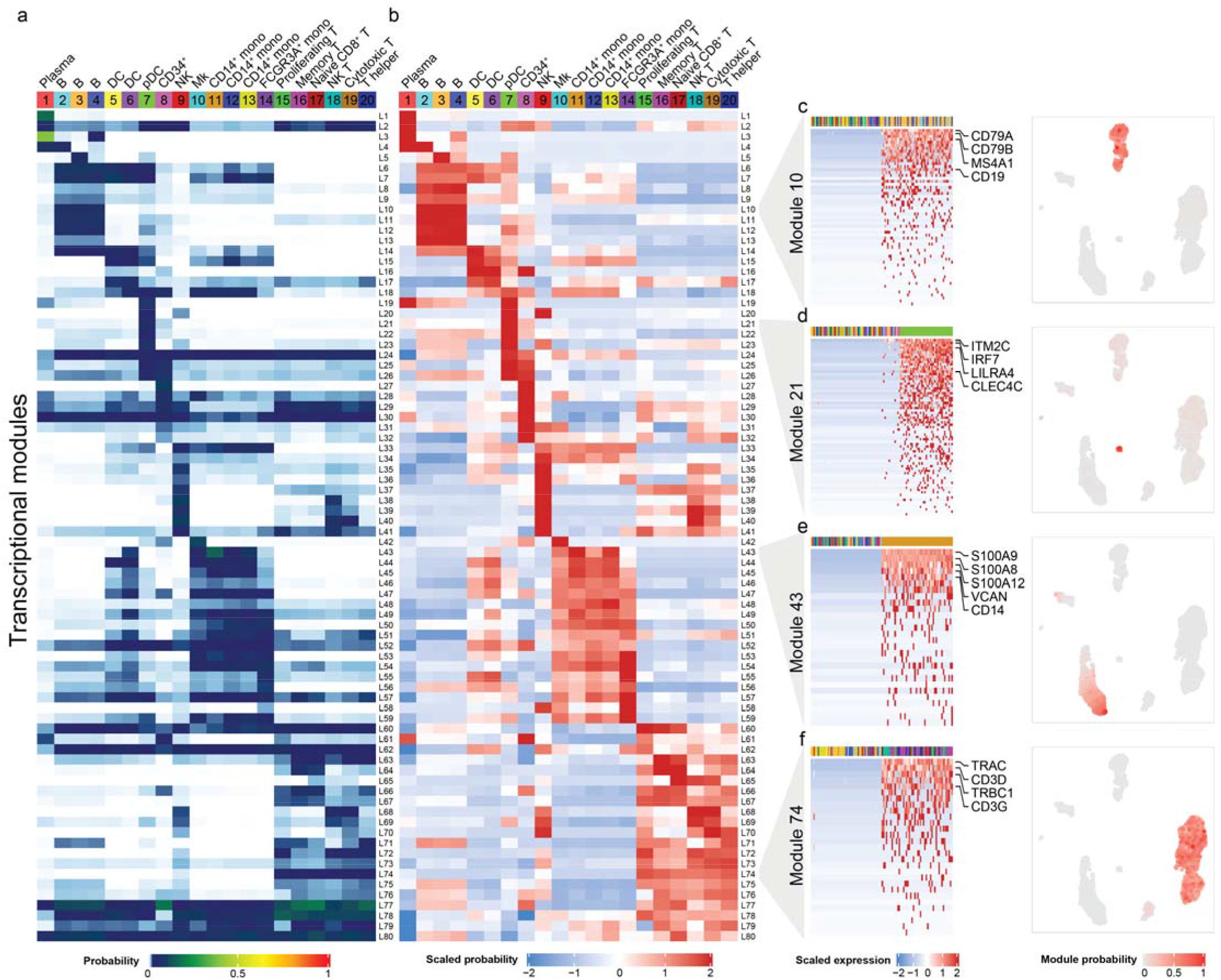
Celda produces a high-level overview of the relationships between transcriptional modules and cell populations. **a**, The *φ* matrix shows the probability of each of the 80 explore the relationship between modules within a cell population. **b**, The row-scaled *φ* matrix can be transcriptional modules (rows) in each of the 20 cellular subpopulations (column) and can be used to used to explore the relative probability of each module across cell populations. **c–f**, Module heatmaps and UMAPs showing the gene expression profiles for cell-type specific transcriptional modules 10, 21, 43, and 74. Top annotation row indicates a total of 100 cells with the highest and lowest probabilities in the module and are colored by their cell cluster labels. Selected marker genes for B cells (CD79A, CD79B, MS4A1, CD19), pDCs (ITM2C, IRF7, LILRA4, CLEC4C), CD14+ monocytes (S100A9, S100A8, S100A12, VCAN, CD14), and T cells (TRAC, CD3D, TRBC1, CD3G) are highlighted on the right.

Traditional single-cell workflows such as those utilized in Seurat^19^ and Scanpy^34^ seek to identify genes that are specific to cell populations using differential expression between that population and all other cells. In Celda, several modules are specific to individual cell populations or cell types. For example, module 10 is expressed in clusters 2, 3, and 4 and contains the B lymphocyte antigen receptor genes CD79A, CD79B as well as the B lymphocyte cell surface antigens MS4A1 and CD19 (**Figure 3c**). Module 21 contains pDC marker genes ITM2C, IRF7, LILRA4, and CLEC4C and has high probability in cell population 7 (**Figure 3d**). Module 43 contains monocyte cell markers S100A9, S100A8, S100A12, VCAN, CD14, and has high probabilities in cell populations 11, 12, and 13 (**Figure 3e**). Modules 74 contains T-cell receptor genes TRAC, CD3D, TRBC1, and CD3G and has high probabilities in cell populations 15-20 (**Figure 3f**). UMAPs colored by module probabilities can illustrate the patterns of transcriptional modules across cell populations.

In addition to the identification of co-expressed genes specific to a single cell type, Celda gene modules can also be used to identify transcriptional programs that are jointly expressed across multiple cell populations. For example, transcriptional modules 12, 44, 40, and 65 have high probability in at least two unique cell subpopulations (**Figure 4a**). Module 12 contains genes BANK1 and BLNK associated with B-cell activation, and genes FCGR2B and HLA-DOB associated with antigen processing and presentation and have high probability in both B-cells and pDCs. Module 44 contains genes including LYZ and SIRPA that are associated with both DCs and CD14+ monocytes. Module 40 is present across NK-cells, cytotoxic T-cells, and NKT cells and contains granzyme genes such as GZMA and GZMH important for cytolytic activity^35^. Module 65 is expressed in both naive and cytotoxic T-cells and contains genes for the CD8 receptor, CD8A and CD8B. Transcriptional modules 15, 45, 47, and 75 are present in at least three unique cell subpopulations (**Figure 4b**). Module 15 contains myeloid lineage genes CD33, CSF2RA, and IL1R2. Module 45 contains toll-like receptor genes TLR2, TLR4, and TLR8. Module 47 contains C-type lectin domain family genes CLEC4A, CLEC7A, CLEC12A, CLEC4G, and leukocyte immunoglobulin-like receptor genes LILRA2 and LILRB3. These three modules all have high probabilities to varying degrees in DCs, pDCs, and monocytes. Module 75 contains lymphoid lineage marker CD69 and has high expression in B-, T-, and NK-cells. Modules 7, 14, 6, and 33 span 4 unique cell subpopulations (**Figure 4c**). Modules 7, 14, and 6 have high probability in B cells, DCs, pDCs, and monocytes. Modules 7 and 14 are predominated by MHC class II genes which are key determinants of antigen presenting cells^36^ (APCs). Module 6 contains CD74 which is an important chaperone that regulates antigen presentation^37^. Module 33 contains genes such as transmembrane immune signaling adaptor TYROBP, IgE receptor gene FCER1G, and macrophage inflammatory gene CCL3 and is expressed in DCs, pDCs, monocytes and NK cells. Modules 24, 28, 62, and 80 have high probability in almost all cell populations and contain many known housekeeping and essential genes. Module 28 contains several common housekeeping genes such as GAPDH, HMGB2, HMGB3, and TUBA1C^38^. Module 80 contains mitochondrial genes MT-CO1, MT-CO2, and MT-CO3. Although expressed to varying degrees in all cells, an extremely high proportion of these genes can indicate severe stress or poor quality within a cell^39, 40^.

**Figure 4.**
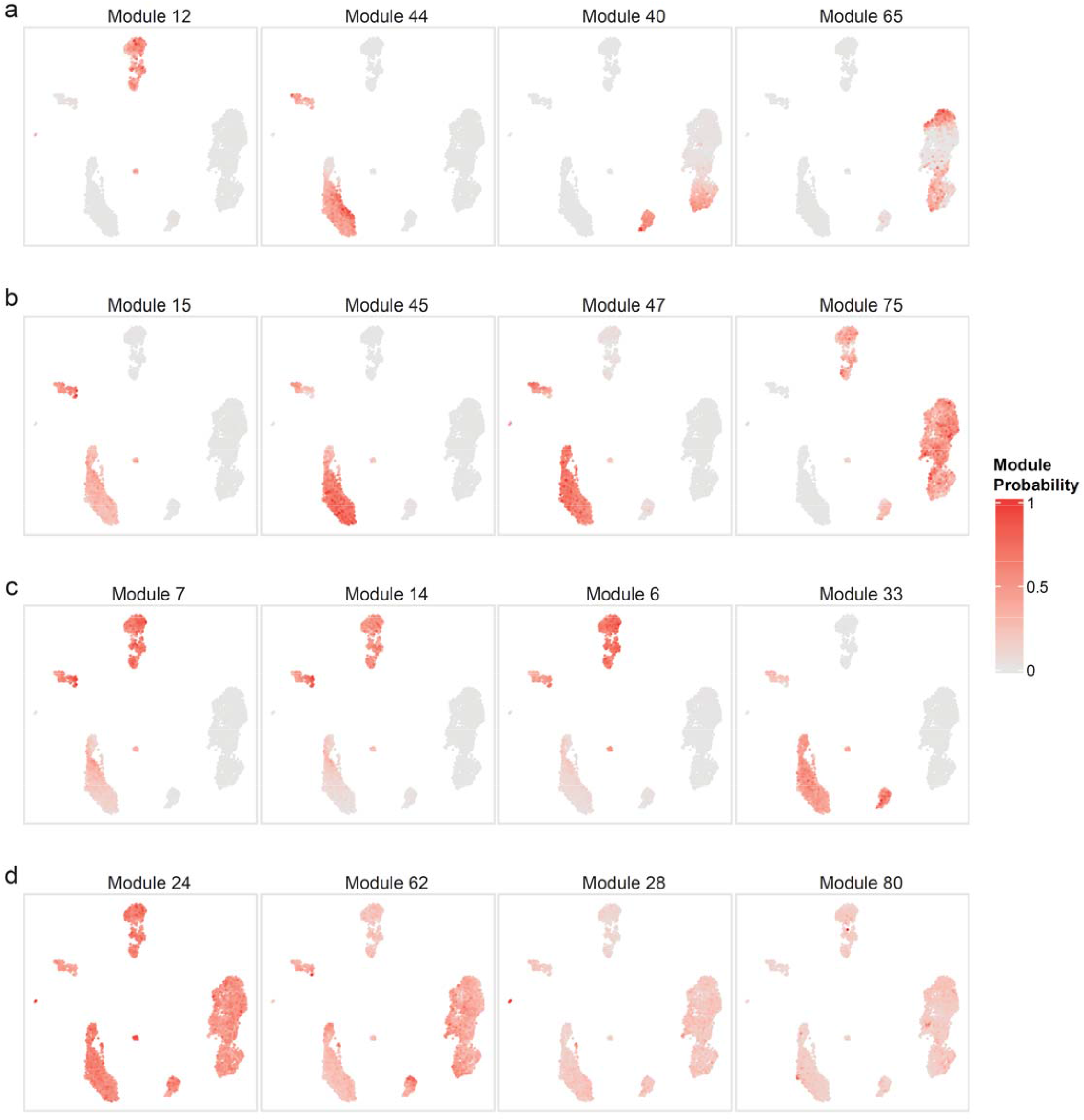
Celda identifies transcriptional modules shared across cell populations. **a**, Selected example UMAPs showing modules with high probabilities in at least two different cell types: module 12 in B cells and pDCs, module 44 in DCs and CD14+ monocytes, module 40 in NK and NK T cells, module 65 in naive cytotoxic T and cytotoxic T cells. **b**, Selected UMAPs showing modules with high probabilities in at least three different cell types: modules 15, 45, and 47 in DCs, pDCs and monocytes, and module 75 in B, T, and NK cells. **c**, Selected UMAPs showing modules with high probabilities in at least four different cell types: modules 7, 14, and 6 in B cells, DCs, pDCs, and monocytes, and module 33 in DCs, pDCs, monocytes, and NK cells. **d**, Selected UMAPs showing modules with high probabilities in all 20 cell clusters. Analyzing modules can reveal novel insights about biological programs active in one or more cell types.

### Comparison of Celda to principal components for module detection

Many scRNA-seq clustering workflows, including ascend^9^, Seurat^19^, and TSCAN^21^, perform dimensionality reduction using principal component analysis (PCA) before cell clustering. Genes that have large loading scores (positive or negative) to a principal component (PC) will be highly correlated with that PC and are often plotted together in a heatmap when assessing the quality of PCs^19, 41^. To qualitatively compare transcriptional modules from Celda to those derived using a PCA-based approach, we analyzed the same PBMC dataset using Seurat (**Figure 5**). There are three major issues when trying to define gene modules using PCA. The first issue is that biological programs from different cell types can be represented at each end of the PC. For example, when examining PC2 from the PCA generated by Seurat, the top 15 genes negatively correlated with PC2 contained B-cell markers CD79A, MS4A1, CD79B, and MHC class II genes, while the top 15 genes positively correlated with PC2 contained T- and NK cell marker genes including TRAC, CD3D, CD7, CTSW, and NKG7 (**Figure 5a**). Similarly, the B-cell and pDC subpopulations were enriched with negative PC2 scores, while the NK cells and a subset of T-cells were enriched with positive PC2 scores (**Figure 5b**). The average expressions of the top 15 genes negatively correlated with PC2 and the top 15 genes positively correlated with PC2 further confirmed enrichment of PC2 associated genes in different cell types (**Figure 5c, d**). The second issue is that transcriptional programs co-expressed in a subset of cell populations can be conflated within the same PC. For example, B-cell marker genes such as CD79A, MS4A1, and CD79B were negatively correlated with PC2 along with MHC-class II genes such as HLA-DRA and HLA-DPA1. While the MHC class II genes are highly expressed in the B-cell populations, they are also highly expressed in the dendritic cell populations where B-cell marker genes are absent. The third issue is that a gene can be highly correlated with many PCs. For example, CST7, NKG7 and GZMA were among the top 15 genes in PCs 2, 3, and 4, while CD7 was among the top 15 genes in PCs 1, 2, and 8 (**Supplementary Figure 5**). Overall, these results illustrate that genes from different cell types and different biological programs can be associated with the same PC and a single gene can be associated with multiple PCs.

**Figure 5.**
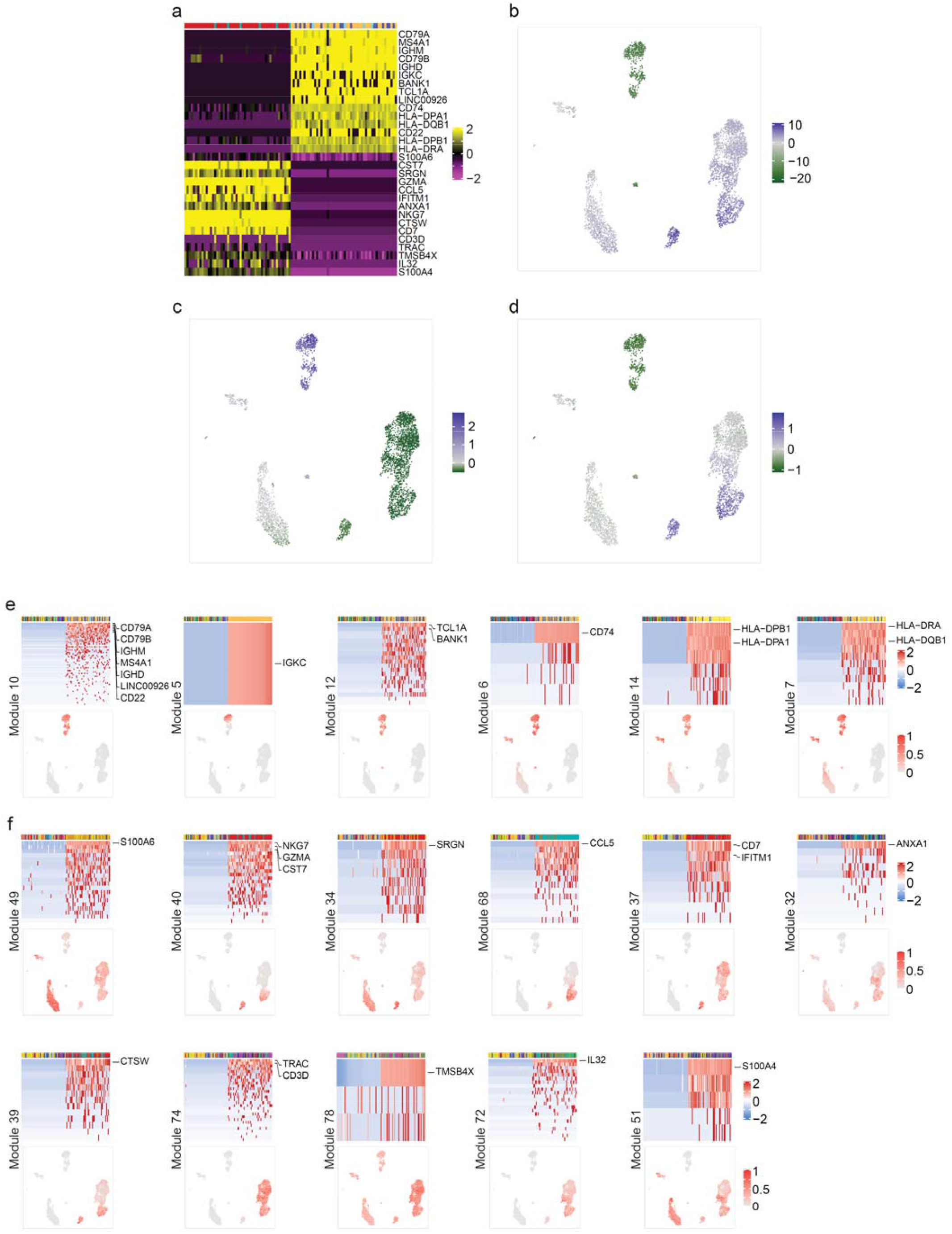
Qualitative comparison of gene co-variation patterns derived from Celda and PCA. **a**, The 15 genes with the most positive loadings for PC2 and 15 genes with the most negative loadings for PC2 are shown in rows of the heatmap. The 50 cells with the lowest PC2 scores and the 50 cells with the highest PC2 scores are showed in the columns of the heatmap. The top annotation row contains Celda cell subpopulation labels. **b**, UMAP colored by scores for PC2. **c**, UMAP colored by the average scaled expression of top 15 genes negatively correlated with PC2. **d**, UMAP colored by the average scaled expression of top 15 genes positively correlated with PC2. **e**, Heatmaps and UMAPs of 6 Celda modules containing the top 15 genes negatively correlated with PC2. **f**, Heatmaps and UMAPs of 11 Celda modules containing the top 15 genes positively correlated with PC2. Overall, these results show that the genes most highly correlated with PC2 from PCA can have different patterns of expression across cell types. In contrast, Celda provided additional insight to gene co-variation by categorizing these top genes into more refined transcriptional modules.

Celda provided additional insight to gene co-variation by categorizing these top genes into more refined transcriptional modules. For example, among the top 15 genes negatively correlated with PC2, 7 genes were in module 10 and expressed across all B-cell subpopulations (**Figure 5e**). However, other genes were clustered in 5 other modules and exhibited different patterns across cell populations. IGKC was found in module 5 by itself and was expressed only in one of the B-cell subpopulations (cell cluster 3). TCLA and BANK1 were in module 12 which was present in B-cell and pDC populations. Similarly, the MHC-class II associated genes were found in modules 6, 14, and 7. These three modules had high probability in B-cell, DC, and pDC populations and moderate probability in different subsets of monocyte populations to varying degrees. Among the top 15 genes positively correlated with PC2, 5 were clustered in modules 40, 68, and 39 (**Figure 5f**). These modules showed enrichment in NK, NKT, and cytotoxic T-cell populations which were enriched with positive PC2 scores. However, 10 remaining genes clustered in 8 other modules showed patterns undetected by PC2. For example, S100 family genes S100A6 and S100A4 were found in modules 49 and 51, which were present in DC, monocyte, and subsets of T- and NK cells. SRGN was in module 34 which was present in NK cells, DCs, pDCs, monocytes, and had moderate probability in T-cells. CD7 and IFITM1 were found in module 37 which had high probability in NK and T-cells. Annexin family gene ANXA1 was grouped in module 32 which was present in subsets of T-cells, NK cells, monocytes, DCs, and CD34^+^ cells to varying degrees. T-cell receptor genes TRAC and CD3D were grouped in module 74 which was present across all T-cell subpopulations. TMSB4X was grouped in module 78 which had high probability in T-cells and moderate probability in all other cell populations. IL32 was in module 72 which had high probability in proliferating T-cells and moderate probability in other T-cell populations. Overall, these results suggest that Celda can identify transcriptional programs representing unique biological processes with better clarity than what can be readily parsed by associating genes with PCs from PCA.

To systematically benchmark Celda’s ability to cluster genes into modules, we compared the performance of Celda_CG and PCA to accurately cluster genes into modules based on simulated data. To create distinct, non-overlapping modules from PCA, each gene was assigned to a single PC based on the magnitude of its loading ranks across all PCs (**Methods**). Six datasets were simulated with increasing similarity between cell populations and modules (**Figure 6**). Celda_CG outperformed PCs in clustering co-expressed genes into transcriptional modules for all simulated clustering difficulties (**Figure 6a**). Median adjusted rand indices (ARIs) for the six increasing clustering difficulties were 0.98, 0.88, 0.84, 0.57, 0.10, and 0.01 for Celda_CG, and 0.20, 0.06, 0.04, 0.02, 0.01, and 0.01 for PCs. These results demonstrate that Celda_CG was more accurate at identifying modules of co-expressed genes compared to a PCA-based approach.

**Figure 6.**
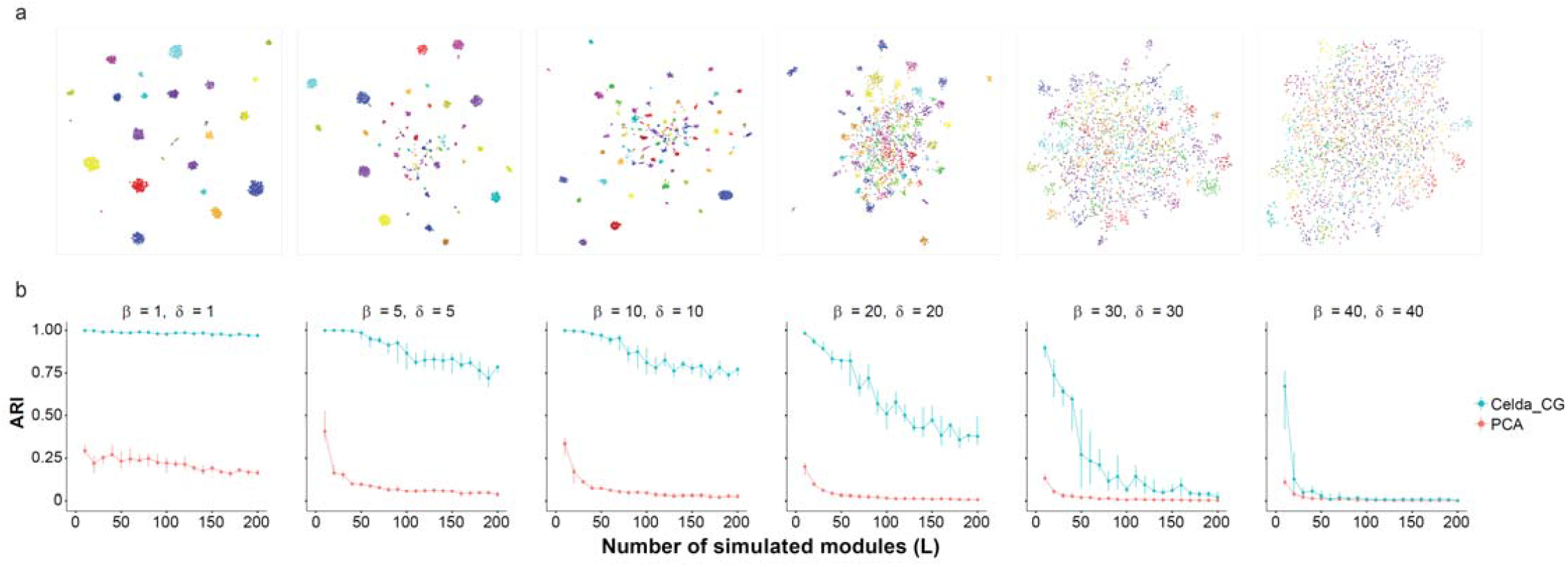
Celda_CG achieves a higher accuracy for clustering of genes into modules compared to a PCA-based approach. Datasets were simulated according to the generative process of the Celda_CG model. Higher values of *β* produced more similar transcriptional modules within each cell population and higher values of *δ* produced a more equal distribution of counts between genes within each module. A range of *L* values from 10 to 200 was simulated for each combination of the parameters. **a**, UMAPs for one of ten replicate simulations at *L = 100* were generated to show the relationship between the 2,000 most variable genes. Each point on the UMAP represents a single gene and is colored by its true module label. Genes closer together in the UMAP have more similar expression patterns across cells. **b**, The adjusted Rand index (ARI) shows the similarity between the true module labels and the gene clustering results for Celda_CG or PCA. Points and vertical lines represent medians and interquartile ranges of ten replicate simulations. Celda_CG achieved a higher median ARI compared to PCA for all *L* values in 5 out of 6 datasets.

## Discussion

Celda is a novel discrete Bayesian hierarchical model for scRNA-seq data that can perform co-clustering of cells into subpopulations and genes into transcriptional modules. When applied to a well-characterized PBMC dataset, Celda revealed novel cell populations missed by other approaches and provided information about the combination of transcriptional programs that distinguished each population. Raw scRNA-seq count data are generally discrete and sparse after UMI corrections are applied. Many available workflows that perform cell clustering for scRNA-seq count data often requires preprocessing the data before clustering. Seurat^19^, ascend^9^, TSCAN^21^, SC3^17^, CIDR^11^, and scran^18^ all perform cell clustering based on dimensionality reduced data, which requires some of the preprocessing steps including normalization of total counts in each cell, logarithmic transformation, and/or z-score standardization to center and scale the variables. Celda is based on hierarchical Dirichlet-multinomial distributions which inherently work with sparse non-negative integer count data without prior normalization. Multinomial distributions have been shown to model UMI-corrected data without inflation better than conventional normalization strategies^42^. For example, the single plasma B-cell was identified by Celda because it had nearly twice the raw counts compared to any other cell in the PBMC dataset. We also used the top 2,000 most variable genes determined by variance-stabilizing transformation^19^ for clustering the PBMCs in this particular analysis. We note that this is not a requirement when running Celda. For example, we previously clustered this dataset with Celda by including 4,529 genes with at least 3 counts across 3 cells^43^. While the overall cluster solutions are similar, applying the variability filter in this analysis promoted the clustering of CD8A and CD8B into a unique module that helped to define the naive CD8^+^ T-cell population. In general, limiting to variable genes can decrease the computational time and help identify modules of genes with lower overall counts but will exclude some genes from being characterized in transcriptional modules.

One major challenge with clustering tools applied to any data type is determining the number of clusters. Statistical metrics to assess cluster stability can be used in conjunction with prior biological knowledge to settle on a solution that is robust and gives the most biological insight. Seurat implements modularity-based community detection where a resolution parameter is used to customize the granularity level at which community structures are detected but does not provide inherent metrics for choosing the number of clusters^19,41,44^. ascend sets a supervised pruning window in the agglomerative hierarchical clustering procedure using Ward’s minimum variance to determine the number of subpopulations^9,45^. TSCAN uses Gaussian mixture modelling which relies RPC^28^ to assist in choosing the number of cell clusters (*K*) and transcriptional modules (*L*). We note on Bayesian information criterion (BIC) to determine the number of clusters^21,46^. In Celda, we use However, further splitting of modules or cell populations by choosing higher *L* or *K* may be useful in that the elbows in the RPC plots can provide good starting point for choosing these numbers. some settings and can be performed after examining UMAPs and module heatmaps. Another limitation of our current model is that technical differences between batches of samples are not taken into account. In the future, we plan to develop specific distributions that can specifically model technical variation between groups of samples. Overall, Celda presents a novel model-based clustering approach towards simultaneously characterizing cellular and transcriptional heterogeneity in biological samples profiled with scRNA-seq assays.

## Methods

### Celda_CG Statistical model

Celda_CG model uses sets of Dirichlet-multinomial distributions to model the hierarchies in the scRNA-seq data. The generative process for celda_CG is outlined in **Figure 1** and below while the complete specification for the model can be found in the **Supplementary Information**. The generative process for Celda_CG is as follows:

1. Draw *η* ∼ Dir _*L*_(*γ*)
2. For each gene *g* ∈ {1,2…, *G*}, draw *y*_*g* ∼_ Mult (*η*)
3. For each transcriptional module distribution *l* ∈ {1,2…, *L*}
  a. Define 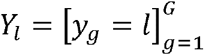
  b. Draw *ψ*_*l*_ ∼Dir (*δY*_*l*_)
4. For each sample *i* ∈ {1,2…, *S*} draw *θ* _*i*_ ∼ Dir _*K*_ (*α*)
5. For each cell Population *k* ∈ {1,2…, *K*}, draw *φ* _*k*_ ∼ Dir _*L*_(*β*)
6. For each cell *j* ∈ {1,2…, *M* _*i*_}, in sample *i*
  a. Draw *Z*,_*i,j*_ ∼ Mult (*θ*_*i*_)
  b. For the *t*-th transcript in cell *j* in sample *i,t* ∈ {1,2…, *N* _*i,j*_},
    i. Draw 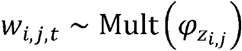
    ii. Draw 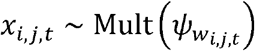

*η* is from a Dirichlet distribution with symmetric concentration *γ* parameter with length equal to the total number of transcriptional modules specified by *L*.*G* is the number of genes. *y*_*g*_ is the hidden transcriptional module label drawn from *η* for gene *g* and will return a value between 1 and *L*.*L* is the number of transcriptional modules. “[]” refers to a Boolean operator and returns 1 when the expression within the bracket is true and 0 otherwise. We use this operator in step 3a to denote that the element corresponding to gene *g* in *Y*_*l*_ will be set to 1 if *y*_*g*_ *= l* and 0 otherwise. *Y*_*l*_ will then be used as an indicator variable in step 3b to control the genes turned on in transcriptional module *l*·*ψ* _*l*_ is from a Dirichlet distribution parameterized by *δY* _*l*_ where each element represents the probability of a gene in the module. If an element in *Y*_*l*_ is zero, the parameter *δY* _*l*_ for the Dirichlet distribution will be zero along with the corresponding probability *ψ*_*l*_ for that gene, thus turning off the expression of that gene in that module. The combination of these variables results in the “hard-clustering” behavior by controlling the assignment of each gene to a single transcriptional module. *S* is the number of samples. *θ*_*i*_ is from a Dirichlet distribution parameterized by the symmetric concentration parameter *α* that defines the probability of each cell population in each sample *i. K* is the number of cellular subpopulations. Each cell population *k* follows a Dirichlet distribution *φ*_*k*_ parameterized by the symmetric concentration parameter *β* where each element in *φ*_*k*_ represents the probability of a transcriptional module in population *k. M*_*i*_ is the number of cells in sample *i*. *Z*_*i,j*_ is the hidden cell population label for cell *j* in sample *i*. *N*_*i,j*_ is the number of transcripts for cell *j* in sample *i*. *w*_*i,j,t*_ □ is the□hidden□transcriptional□module label□for□transcript□*x*_*i,j,t*_, and □*x*_*i,j,t*_ is □ the □ *t*-th transcript for cell *j* in sample *i* . *z*_*ij*_ is drawn from *θ*_*i*_ and represents the hidden label denoting the population assignment for each cell. *W*_*i,j,t*_ is the hidden label for transcript *t* in cell *j* drawn from 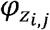 and represents the module assignment for that transcript. *x*_*i,j,t*_ is the observed transcript which is drawn from 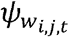. Note that only the genes “turned on” according to the indicators *Y*_*l*_ will have a non-zero probability and will be selected from this draw.

The complete likelihood function of Celda_CG model is then given as:

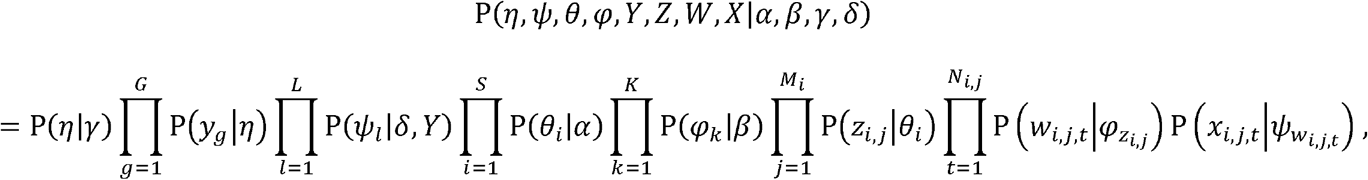

where *α, β, γ, δ* are the symmetric prior parameters in their corresponding Dirichlet distributions, and *Y, Z, W, X*, are the collections of *y*_*g*_, *z*_*ij*,_ *w*_*i,j,t*,_ *x*_*ijt*_ respectively.

### Estimation of model parameters

We use a heuristic hard Expectation Maximization (EM) procedure to estimate the cell population label *z*_*ij*_ for cell *j* in sample *i* and a collapsed Gibbs sampling procedure to estimate the hidden transcriptional module label *y*_*g*_ for gene *g* (**Supplementary Information**). To estimate the hidden transcriptional module label for each gene, we integrate out *ψ, φ*, and *W* and drop components related to *θ* that are invariant with respect to *Y*. The final formula after simplification is as follows:

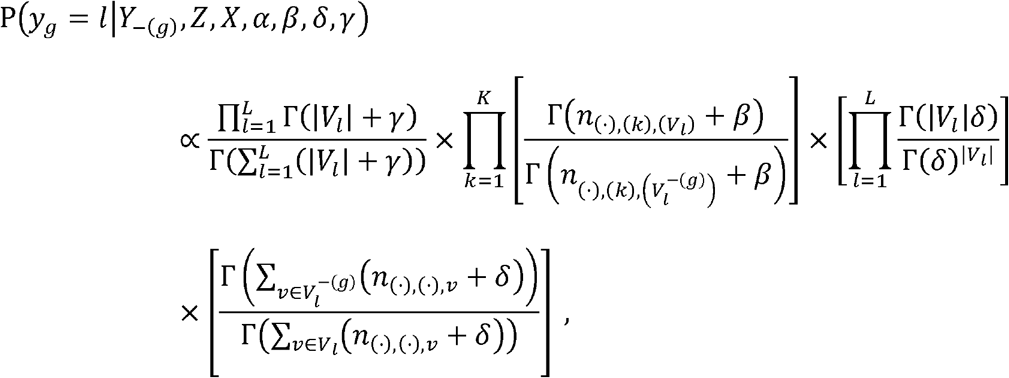

where *L* is the total number of modules, *K* is the total number of cell populations, |*V*_*l*_| is the total number of genes in module 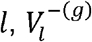 is the total number of genes in module *l* leaving out current gene 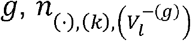 is the total number of transcripts from genes in module *l* cross all the cells in cluster *k* leaving out those from gene *g,n*_(·),(·),*v*_ is the total number of transcripts for gene *g* across all cells and samples, 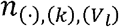 is the number of transcripts from all genes in module *l* in population *k*, and Γ is the gamma function. For estimating the hidden population label for each cell *z*_*i,j*_, we relied on a heuristic “hard” EM procedure to increase speed on large datasets with many cells. The collapsed Gibbs sampling equations for *z*_*i,j*_, can also be found in the **Supplementary Information**. First, we drop components related to ψ that are invariant with respect to *Z*. The “hard” EM obtains a point estimate of *z*_*i,j*_ by maximizing the posterior with respect to point estimates of *θ* and *φ* given the current configurations of *Z* and *Y*:

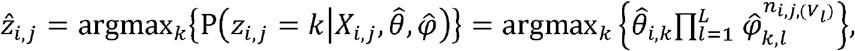

where *X*_*i*_,_*j*_ is the collection of transcripts *x* _*i*_,_*j*_,_*t*_ within cell *j* in sample 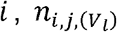 is the number of transcripts from all the genes that belong to module *l* of cell *j* in sample *i*. 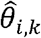 is the point estimate of, *θ*_*i,k*_ which is the probability of a cell belonging to cell population *k* in sample *i* and can be calculated as:

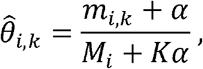

where *m*_*i,k*_ is the total number of cells assigned to cluster *k* in sample *i* and *M*_*i*_ is the total number of cells in sample *i*.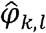 is the point estimate of *φ*_*k,l*_ which is the probability of module *l* in cell population *k* and can be calculated as:

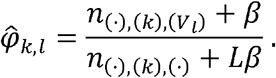

Within each iteration of the optimization procedure, we apply the “hard” EM procedure to estimate the cell population labels (*Z*) given a fixed set of transcriptional module labels (*Y*) and then apply the collapsed Gibbs sampling procedure to estimate the transcriptional module labels (*Y*) given a fixed set of cell population labels (*Y*). We generally run the model for a maximum number of iterations (200 by default) or until there has been no improvement in the log-likelihood for a predefined number of iterations (10 by default). The configuration of *Z* and *Y* that produced the highest likelihood are returned as the final solution.

In order to avoid local optimum, we apply a heuristic cluster/module splitting procedure every 10 iterations. To apply the cell splitting procedure at a given iteration, we try to find a better configuration for *Z* with a higher log-likelihood by splitting one population into 2 new clusters and removing another unsplit population. Let *K*\* be the set of cell population clusters that have more than 3 cells and |*K*\*| be the cardinality of set *K*\*. For one cell population *k*\*∈ *K*\*, the cluster is split into 2 new clusters using the Celda_C model setting *K*_*c*_ = 2. Then parallelly for all other cell clusters {*k*’: *k*’ ∈ {1,2,…;, *K*}∧ *k*’ ≠ *k*\*}, we redistribute all the cells in cluster *k*’ to their second most likely cluster according to to EM probabilities of current *Z* configuration. The log-likelihood is re-calculated for each of these *K* − 1 configurations. After repeating this procedure for all the {*k*\*: *k*\* ∈ *K*\*}, a total number of |*K*\*| × (*K* − 1) new possible configurations for *Z* are obtained. The configuration that produced the highest likelihood will be set at the current solution. If none of the new configurations had a higher likelihood than the original configurations, then no splitting will be performed and the original configuration of will be maintained. The module splitting procedure is similarly applied to the transcriptional modules to find a *Y* that has higher log-likelihood. One module *l*^*^. is split using Celda_G with *L*_*G*_ = 2 and new likelihoods are calculated by redistributing the genes in each of the other modules. One potential limitation is that running Celda_G on all cells to split each module would result in dramatic reduction in speed for large datasets. We therefore take each cell population cluster and split it up into 10 new clusters using Celda_C with *K* = 10 to produce a temporary configuration denoted *Z*^*^. These temporary populations are used to potentially find a better configuration of module labels. Splitting each cell population into 10 temporary cell populations ensures that better splits of the modules can be obtained even if the current cluster labels *Z* are suboptimal. Even though the modules are split with *Z*^*^, the overall likelihood for all new splits of *Y* is still calculated with the current configuration of containing of *Z* containing the *K* subpopulations. As in the cell splitting approach, the module split with the best log-likelihood is chosen if it is higher than that from the current *Y* configuration.

### Determining the number of cell populations and transcriptional modules

Perplexity has been commonly used in the topic models to measure how well a probabilistic model predicts observed samples^5^. Here, we use perplexity to calculate the probability of observing expression counts given an estimated Celda_CG model. Rather than performing cross-validation which is computationally expensive, a series of test sets are created by sampling the counts from each cell according to a multinomial distribution defined by dividing the counts for each gene in the cell by the total number of counts for that cell. Perplexity is then calculated on each test set with a lower perplexity indicating a better model fit^5^. For a test set *x*, the perplexity of Celda_CG is given as

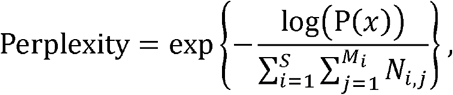

where log(P(*x*)) is defined as:

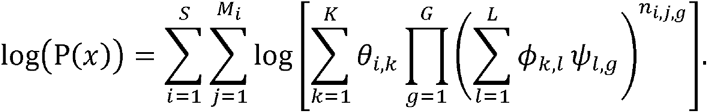

We compare perplexity values among different model settings and use rate of perplexity change^28^ (RPC) to determine an appropriate number of cell populations and transcriptional modules. sequence of equally spaced *K* s arranged in ascending order are fitted. We then calculate the RPC Particularly, setting a fixed number of transcriptional modules, a series of Celda_CG models with a along the course of increase of cell populations, choose the smallest *K* as the appropriate number of cell populations where the RPC is zero at a given precision. Similarly, setting a fixed number of cell populations, an appropriate number of transcriptional modules can be selected by calculating the RPC along a sequence of equally spaced *L* s.

### Data collection and preprocessing

PMBC_4K dataset was downloaded using R/Bioconductor package `TENxPBMCData` v1.8.0. It contains 4,340 cells and 33,694 genes. We applied DecontX^43^ to remove inadvertent contamination using default settings. 17,039 genes detected in fewer than 3 cells were excluded. We applied `NormalizeData` and `FindVariableFeatures` functions from `Seurat` v3.2.2^19^ using default settings and identified a set of 2,000 most variable genes for clustering by variance-stabilizing transformation^19^ (VST). PCA was performed on scaled normalized gene expressions using `RunPCA` function from `Seurat` v3.2.2 in default settings. For coloring of UMAPs and module heatmaps, the decontaminated counts were normalized by library size, square-root transformed, centered, and scaled to unit variance. Values greater than 2 or less than −2 were trimmed.

### Selecting the number of transcriptional modules (*L*) and cell clusters (*K*)

We applied two stepwise splitting procedures as implemented in the `recursiveSplitModule` and `recursiveSplitCell` functions in Celda to determine the optimal *L* and *K*. `recursiveSplitModule` uses the celda_G model to cluster genes into modules for a range of possible *L* values between 10 to 200. The module labels of the previous model with *L* − 1 modules are used as the initial values in the current model with *L* modules. The best split of an existing module, evaluated by best overall likelihood, is found to create the *L*-th module. The rate of perplexity change (RPC) was calculated for each successive model generation. For the PBMC 4K dataset, we found the model with 80 transcriptional modules had low RPC and included both known and novel gene programs. m`recursiveSplitCell` uses the Celda_CG model to cluster cells into cell clusters for a range of possible *K* values between 3 to 30. The module labels of genes from model *L* = 80 was used to initialize the modules in `recursiveSplitCell`. We found the model with 20 cell clusters had low RPC and included both known and novel cell populations (**Supplementary Figure 1**). The final Celda_CG model used in this analysis of the PBMC 4K dataset was extracted from the stepwise splitting results using the `subsetCeldaList` function.

### UMAP of PBMC cells based on Celda transcriptional modules

Dimensionality reduction for visualization by Uniform Manifold Approximation and Projection^29^ (UMAP) is performed using the square root transformed module probability (MP) matrix which contains the probability of each transcriptional module in each cell. Specifically, the MP matrix is defined as

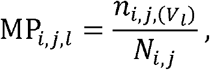

where 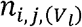 denotes the sum of all counts belonging to genes in transcriptional module *l* and *N*_*i,j*_ is the total sum of counts for a cell. The square root transformation is applied as it can be applied to zero counts without need to add a pseudocount as is required with the log transformation. The `umap` function from the `uwot` R package was applied to the MP matrix using Euclidean distance to obtain 2 dimensional coordinates for each cell with `n_neighbors = 10`, `min_dist = 0.5`, and default settings^29^.

### Testing for differential expression

A hurdle model from MAST^47^ was used for significance testing of differential expression between cell clusters. Benjamini and Hochberg false discovery rate^48^ (FDR) adjusted *p*-values were used to reject the null hypotheses.

### Cell clustering and UMAP using Seurat

The same set of 2000 most variable genes in the decontaminated PMBC dataset were used for clustering by Seurat. PCA was performed on scaled normalized gene expressions using `RunPCA` function from `Seurat`^19^ v3.2.2 in default settings. Shared Nearest Neighbor (SNN) graph was constructed using `FindNeighbors` function with the top 22 PCs. Clusters were identified by modularity optimization using `FindClusters` with default settings. UMAP was generated using `RunUMAP` function and the top 22 PCs with `n.neighbors = 10`, `min.dist = 0.5`, and default settings.

### Cell clustering and UMAP using scran

The same set of 2,000 most variable genes in the decontaminated PMBC dataset were used for clustering by scran. PCA was performed on normalized gene expressions using `runPCA` function from scater^49^ v1.18.3 using 2,000 genes and default settings. SNN graph was constructed using `buildSNNGraph` function from scran^18^ v1.18.3 with the top 28 PCs. Clusters were identified by random walks using `cluster_walktrap` function from `igraph`^50^ package v1.2.6 with default settings. UMAP was generated using `runUMAP` function with `n_neighbors = 5`, and default settings.

### Evaluation of module clustering accuracy with simulated data

Data were simulated based on the generative model of Celda_CG (**Methods** and **Supplementary Information**) using the `simulateCells` function in Celda. Specifically, we set `model` to `“celda_CG”`, `S` to 1, `CRange` between 4,000 and 6,000, `NRange` between 1,000 and 10,000, `G` to 33,000, and `K` to 20. scRNA-seq count data were simulated using six combinations of concentration parameters *β* and *δ* ranging from 1 to 40 representing six levels of clustering difficulties. For each combination of *β* and *δ* a range of the number of transcriptional modules (*L*) from 10 to 200 was simulated with 10 replicates per *L*. After simulated data were generated, the 2,000 most variable genes determined by VST were selected. Celda_CG clustering and PCA were applied to group the 2,000 most variable genes to transcriptional modules and PCs whose numbers equal the true remaining number of modules for the 2,000 genes. Adjusted Rand indices (ARIs) were calculated between the gene clustering results of Celda_CG or PCA and the true module labels of genes using `adjustedRandIndex` function from `mclust` package v5.4.6.

We applied a heuristic approach to cluster genes to PCs based on gene loadings. After performing PCA to reduce the data to PCs whose number equals the number of true modules for the 2,000 most variable genes, we first order the genes by loadings in increasing order for each PC. For each PC, if the sum of the absolute values of the top 50 negative loadings is greater than the sum of the absolute values of the bottom 50 positive loadings, we rank the genes by loadings in this PC in increasing order. Otherwise, we rank the genes by loadings in this PC in decreasing order. This is to account for the bidirectionality of PCA loadings so that when genes are assigned to a PC, they are always in the same direction with respect to the orientation of the PC. After ranking the genes by loadings for each PC, we assign each gene to its highest-ranking PC accordingly. If a gene has the same highest ranks in two or more PCs, this gene is not used for the calculation of ARI.

UMAPs of genes were generated to visualize the variability of genes and the clustering difficulties for the simulated data. For each combination of *β* and *δ*, UMAP was generated based on one of the 10 simulations at *L*=100. Gene counts for each cell were collapsed to cell clusters before applying UMAP. Specifically, for each gene, the counts for each of the 20 true cell clusters were added and divided by total counts for this gene, so the number of features were reduced from the total number of cells to 20. These cell cluster probabilities were then square-root transformed before applying UMAP. UMAP dimension reduction coordinates for cells were generated using the `umap` function from `uwot` R package with `n_neighbors = 10`, `min_dist = 0.5`, and default settings.

## Data availability

The PBMC dataset used in this study is available at https://support.10xgenomics.com/single-cell-gene-expression/datasets/2.1.0/pbmc4k.

## Code availability

The source code for Celda, in the form of an installable R package, is available at the Bioconductor repository https://www.bioconductor.org/packages/celda. The development version is located on GitHub at: https://github.com/campbio/celda. Scripts for reproducing the published results are available at https://github.com/campbio/Manuscripts/tree/master/Celda.

## Acknowledgements

We thank Jiangyuan Liu for contribution of initial differential expression and heatmap code and Paola Sebastiani for reviewing the statistical models. This work was funded by National Library of Medicine (NLM) R01LM013154-01 (JDC, MY) and Informatics Technology for Cancer Research (ITCR) 1U01 CA220413-01 (WEJ, JDC).

## Author Contributions

J.D.C., S.Y., M.Y. and W.E.J. developed Celda statistical model. Z.W., S.Y., Y.K, S.E.C., and J.D.C. implemented the Celda R package. Z.W., Y.K., S.E.C., and J.D.C. performed data analysis. Z.W., S.Y, and J.D.C. wrote the manuscript.

---

**Supplementary Figure 1.**
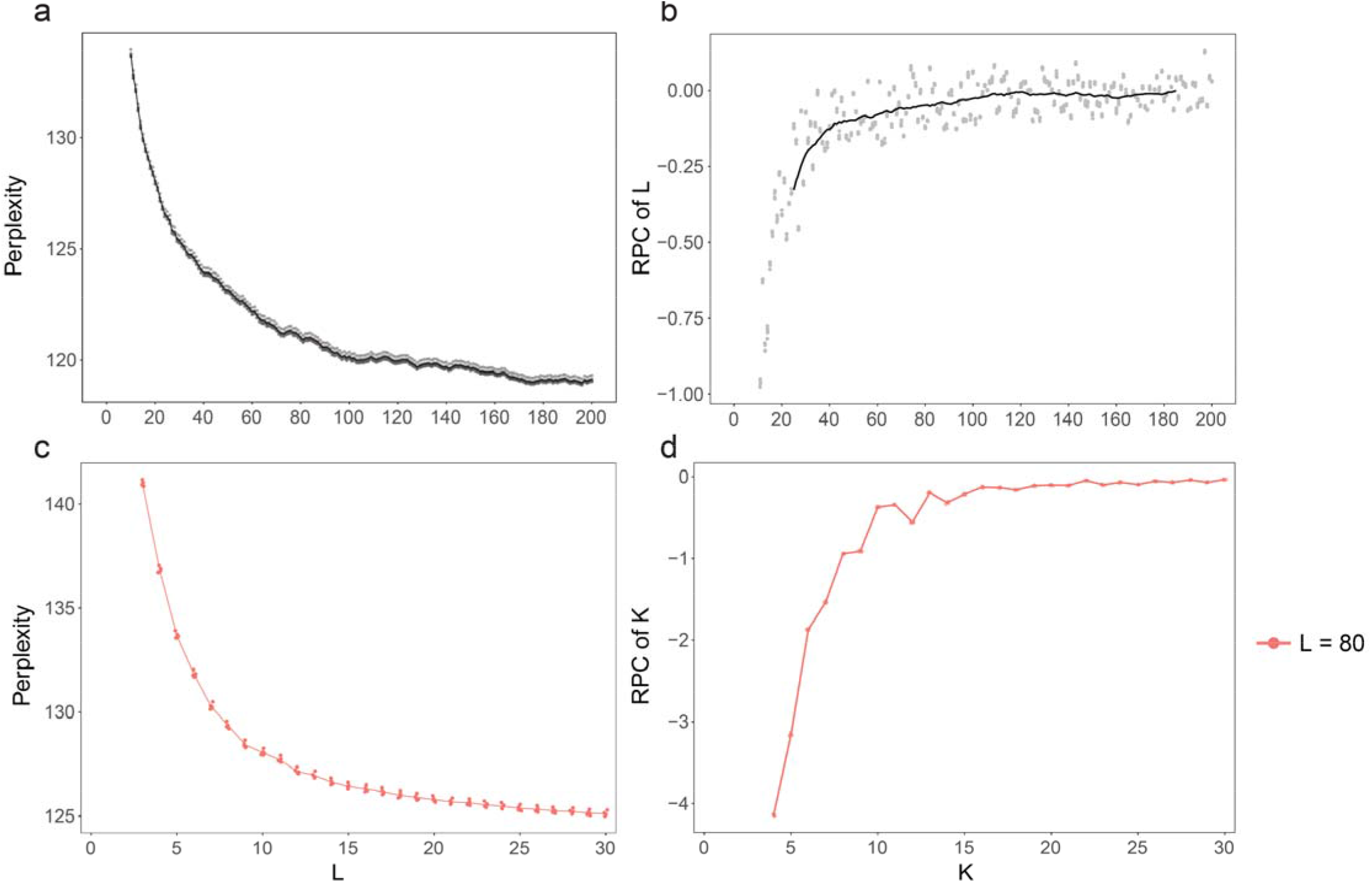
Determining the optimal number of transcriptional modules (*L*) and cell clusters (*K*) for the PBMC 4K dataset. **a**, Scatter plot showing the perplexity of models with a range of 10 to 200 transcriptional modules. **b**, Scatter plot showing the rate of perplexity change (RPC) between each model with *L* transcriptional modules and the previous model with *L* − 1 transcriptional modules. The solid black line represents moving average of centered rolling windows of size 30. For the PBMC 4K dataset, a *L* value of 80 was selected because it was beyond the “elbow” on the curve and captured biologically relevant modules observed when performing manual review of module heatmaps. **c**, Scatter plot showing the perplexity of models with a range of 3 to 30 cell clusters with *L* fixed at 80. **d**, Scatter plot showing the RPC between each model with *K* cell clusters and the previous model with *K* – 1 cell clusters. For the PBMC 4K dataset, a *K* value of 20 was selected because it was beyond the “elbow” on the curve and captured known and novel cell types.

**Supplementary Figure 2.**
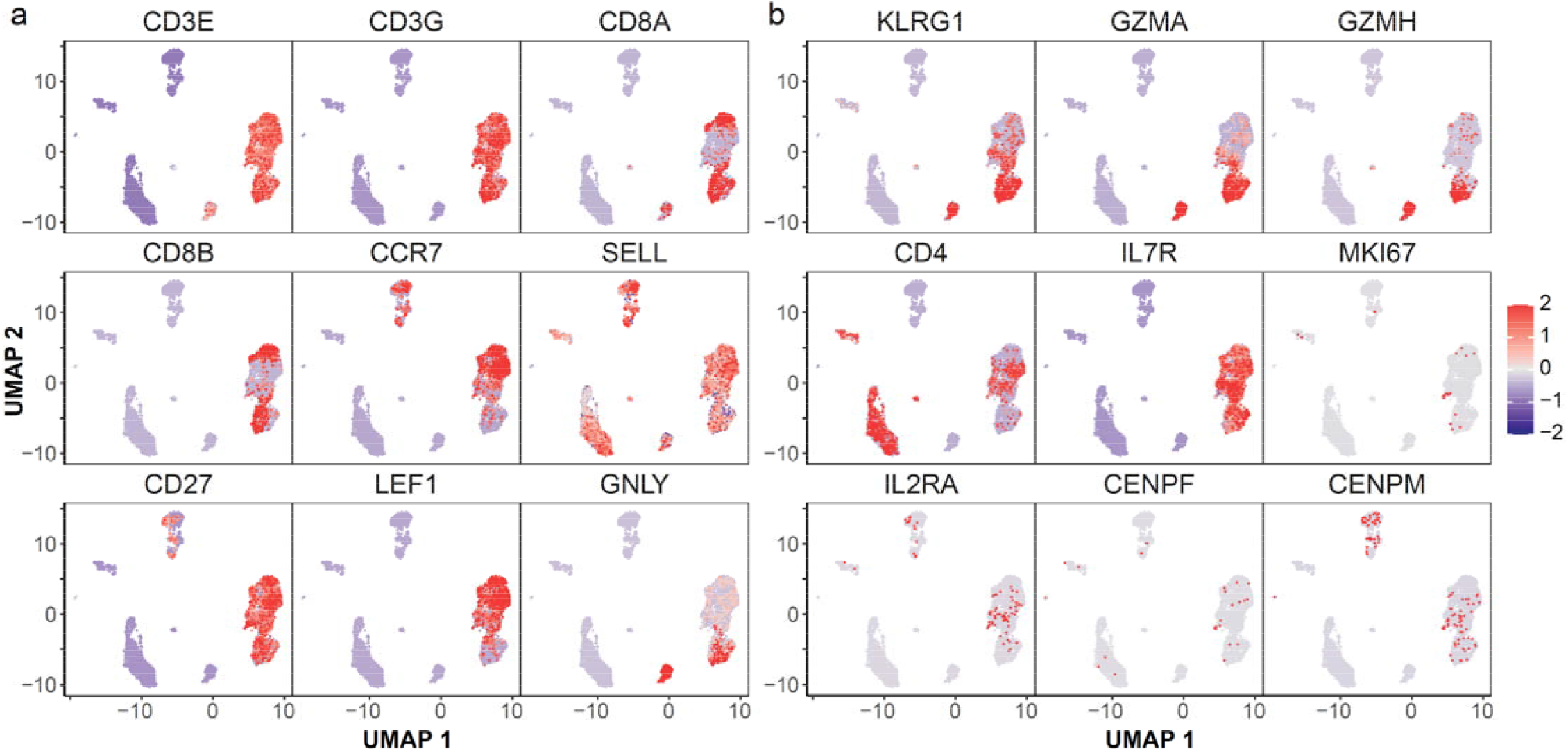
UMAPs showing expression of marker genes for T cell subpopulations. **a**, UMAPs colored by scaled normalized expressions of T cell markers CD3E, CD3G, cytotoxic T cell markers CD8A, CD8B, naive T cell markers CCR7, SELL, CD27, LEF1, and NK T cell marker GNLY. **b**, UMAPs colored by scaled normalized expressions of NK T cell markers KLRG1, GZMA, GZMH, T helper cell markers CD4, IL7R, and activated T cell markers MKI67, IL2RA, CENPF, CENPM.

**Supplementary Figure 3.**
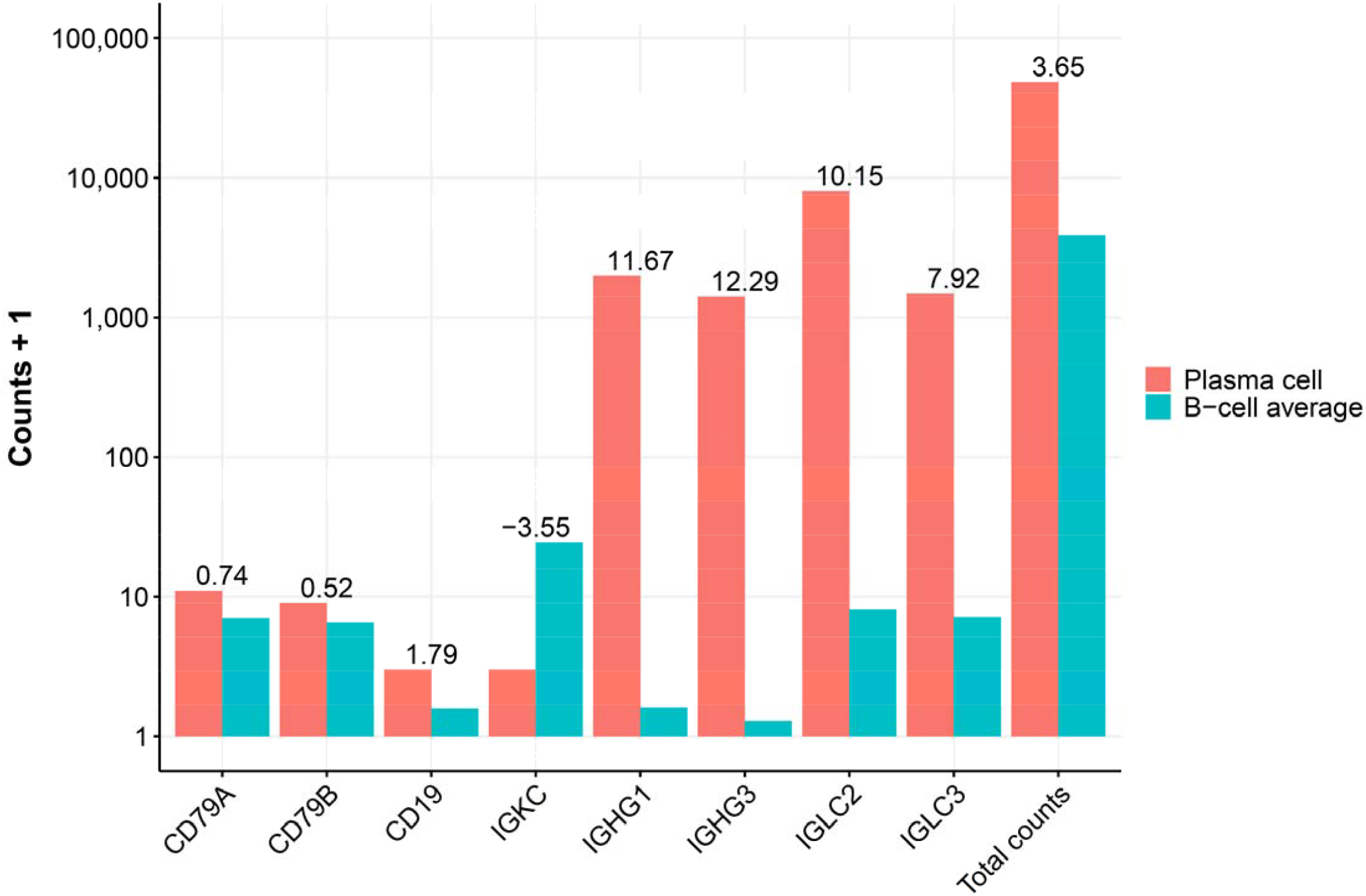
The plasma cell showed much higher expression of IGHG1, IGHG3, IGLC2, and IGLC3 compared to B-cells. The total raw UMI counts (with one pseudocount) of CD79A, CD79B, CD19, IGKC, IGHG1, IGHG3, IGLC2, and IGLC3 for the plasma cell (cell cluster 1) and B-cells (cell clusters 2, 3, 4) are shown on the bar plots. The log_2_ fold change values are shown on the top of bars. The plasma cell showed much higher expression levels of IGHG1, IGHG3, IGLC2, and IGLC3 while having relatively similar expression levels of CD79A, CD79B, and CD19 compared to the average of B-cells.

**Supplementary Figure 4.**
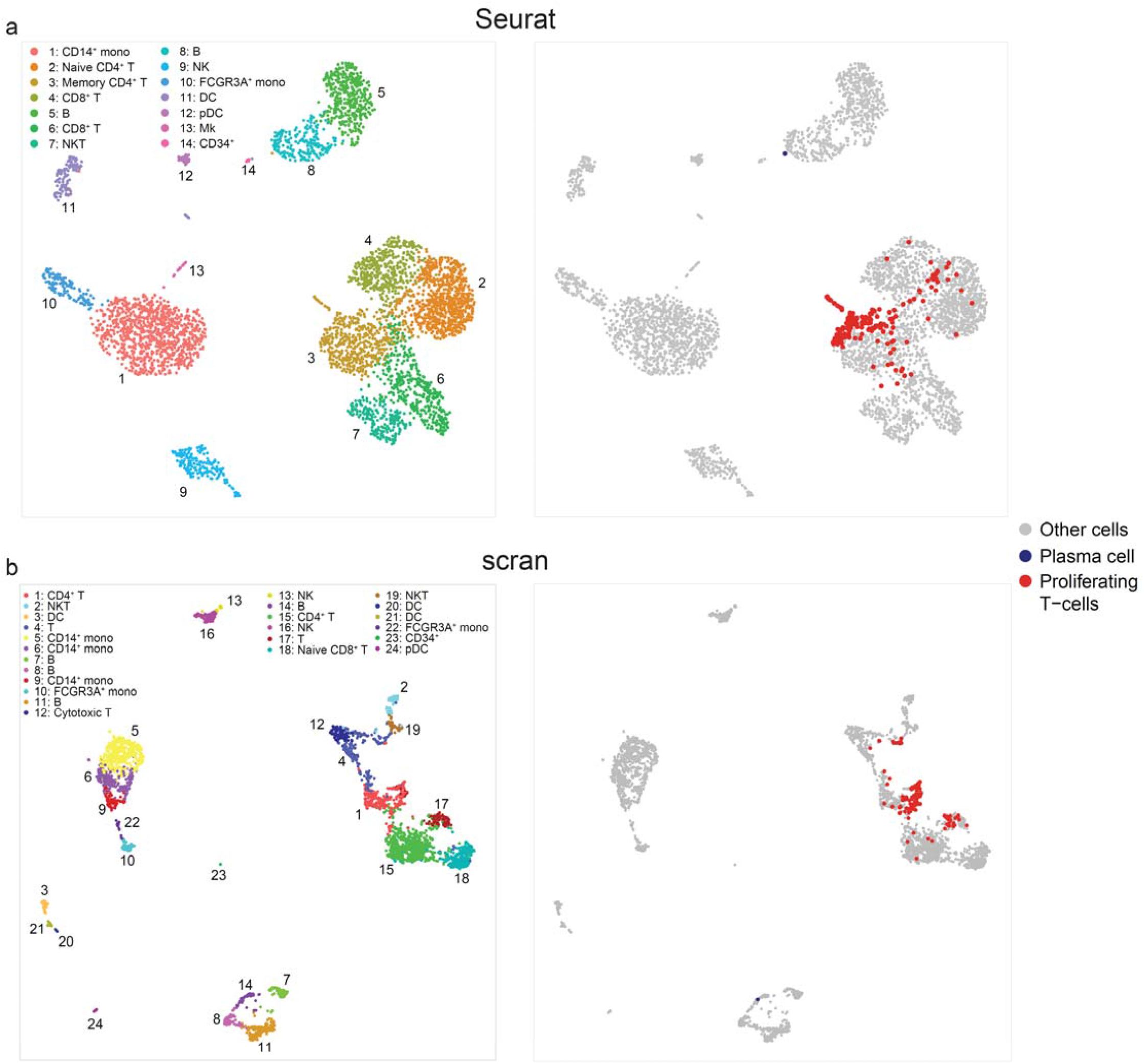
The Proliferating T-cells and plasma cell are not distinctly clustered by Seurat and scran. UMAPs of PBMC cells colored by cell populations identified by Seurat and scran are shown. The proliferating T-cells and plasma cell identified by Celda are highlighted on the right. **a**. Seurat grouped both the proliferating T-cells and the plasma cell in the CD4^+^ T-cell cluster. **b**. scran grouped the proliferating T-cells in the CD4^+^ T-cell cluster and clustered the plasma cell with B-cells.

**Supplementary Figure 5.**
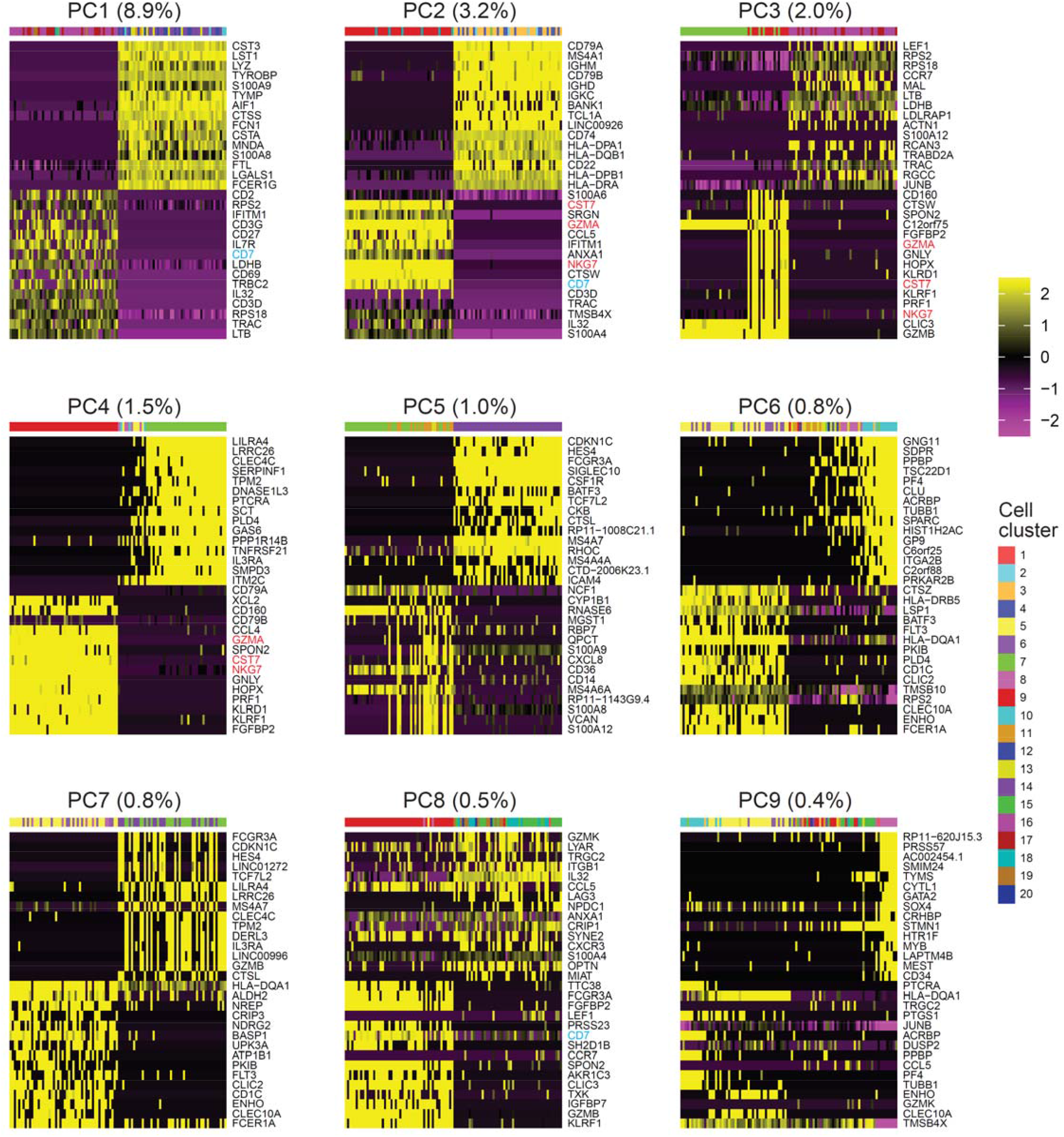
Genes can be highly correlated with many PCs from PCA. Heatmaps of the first 9 PCs colored by scaled normalized gene expressions. Genes are ranked by their loadings in increasing order. The top and bottom 15 genes for each PC are shown. Top annotation row indicates a total of 100 cells with the highest and lowest PC scores and are colored by Celda cell cluster labels as in figures 2 and 3. CST7, NKG7 and GZMA (highlighted in red) are present in the top genes in PCs 2, 3, and 4. CD7 (highlighted in cyan) is present in the top genes in PCs 1, 2, and 8.

**Supplementary Table 1.** Marker genes used to identify cell types in the PBMC dataset.

| Cell type | Celda_CG cell cluster | Marker genes |
| --- | --- | --- |
| Plasma cell | 1 | IGHG1, IGHG3, IGLC2, IGLC3 |
| B cell | 2, 3, 4 | CD19, MS4A1, CD79A, CD79B |
| Dendritic cell | 5, 6 | FCER1A, CLEC10A, FLT3, HLA-DPB1, HLA-DPA1, HLA-DQA1 |
| Plasmacytoid dendritic cell | 7 | ITM2C, IRF7, IRF8, LILRA4, CLEC4C |
| CD34+ progenitor cell | 8 | CD34, SOX4, MYB, GATA2 |
| Natural killer cell | 9 | NKG7, KLRD1, CST7 |
| Megakaryocyte | 10 | PPBP, ITGA2B, PF4 |
| CD14+ monocyte | 11, 12, 13 | CD14, S100A9, S100A8 |
| FCGR3A+ monocyte | 14 | FCGR3A, LST1, SERPINA1 |
| Proliferating T cell | 15 | MKI67, IL2RA, CENPF, CENPM |
| Memory T cell | 16 | CCR7, SELL, CD27 |
| Naive CD8<sup>+</sup> T cell | 17 | CCR7, CD3D, CD8A, CD8B |
| Natural killer T cell | 18 | CD3D, GNLY, KLRG1, GZMA, GZMH, |
| Cytotoxic T cell | 19 | CD3D, CD8A, CD8B |
| T helper cell | 20 | CD3D, CD4, IL7R |

