## Supplementary Information for "Celda: A Bayesian model to perform co-clustering of genes into modules and cells into subpopulations using single-cell RNA-seq data"

### 1 Celda_C: Clustering cells into subpopulations across samples

#### 1.1 Background

The overall goal of the **Celda_C** is to cluster cells with similar count distributions into the same cell population and is similar to previous document clustering models such as the Dirichlet Multinomial Mixture Model [1] or the single-cell clustering method DIMM-SC [2]. However, **Celda_C** also allows for cells from multiple samples to be clustered together and assume that each sample may contain different proportions of each cell population. **Celda_C** will detemine the hidden label for each cell (i.e. cluster assignment to a population), estimate the contribution of each gene in each cell population, and quantify the distrubiton of each cell population for each sample (if multiple samples are available).

We breifly review several properties of the Dirchlet-multinomial distribution used throughout the collapsed Gibbs samplers. Let $\theta$ follow a symmetric Dirichlet distribution of length $K$ parameterized by $\alpha$, that is $\theta \sim \text{Dir}_K(\alpha)$. Here, $\alpha$ is used as a single scalar value that is repeated $K$ times for form the vector $\boldsymbol{\alpha}$, the parameter for the symmetric Dirichlet distribution. The probability density function for this distribution is defined as:

$$p(\theta|\alpha) = \frac{\Gamma(\sum_{k=1}^{K} \alpha_k)}{\prod_{k=1}^{K} \Gamma(\alpha_k)} \prod_{k=1}^{K} \theta_k^{\alpha_k - 1}. \tag{1}$$

Since we will utilize a symmetric Dirichlet distribution where all values of $\alpha_k$ are identical, we can simplify this to:

$$p(\theta|\alpha) = \frac{\Gamma(K\alpha)}{\Gamma(\alpha)^K} \prod_{k=1}^{K} \theta_k^{\alpha - 1}. \tag{2}$$

Let $\boldsymbol{Z}$ be a vector representing a series of $M$ independent draws from a multinomial distribution parameterized by $\theta$, that is $z_i \sim \text{Mult}(\theta)$ for $i = 1..M$. The joint probability distribution of $\boldsymbol{Z}$ is:

$$\prod_{i=1}^{M} p(z_i|\theta) = \prod_{k=1}^{K} \theta_k^{m_k}, \tag{3}$$

where $m_k$ represents the number of items in $\boldsymbol{Z}$ that are assigned to component $k$, that is the number of times $z_i = k$ in vector $\boldsymbol{Z}$. Using the Dirichlet distribution as the prior for the series of multinomial random variables, we can write the joint distribution as:

$$\begin{aligned} p(\theta, \boldsymbol{Z}|\alpha) &= p(\theta|\alpha) \prod_{i=1}^{M} p(z_i|\theta) \\ &= \frac{\Gamma(K\alpha)}{\Gamma(\alpha)^K} \prod_{k=1}^{K} \theta_k^{\alpha - 1} \prod_{k=1}^{K} \theta_k^{m_k} \\ &= \frac{\Gamma(K\alpha)}{\Gamma(\alpha)^K} \prod_{k=1}^{K} \theta_k^{m_k + \alpha - 1}. \end{aligned} \tag{4}$$

To build a collapsed Gibbs sampler, we can integrate out $\theta$ as follows:

$$\begin{aligned}
p(\boldsymbol{Z}|\alpha) &= \int_\theta p(\theta, \boldsymbol{Z}|\alpha) d\theta \\
&= \int_\theta p(\theta|\alpha) \prod_{i=1}^{M} p(z_i|\theta) d\theta \\
&= \int_\theta \frac{\Gamma(K\alpha)}{\Gamma(\alpha)^K} \prod_{k=1}^{K} \theta_k^{m_k+\alpha-1} d\theta \\
&= \frac{\Gamma(K\alpha)}{\Gamma(\alpha)^K} \frac{\prod_{k=1}^{K} \Gamma(m_k+\alpha)}{\Gamma(\sum_{k=1}^{K}(m_k+\alpha))} \int_\theta \frac{\Gamma(\sum_{k=1}^{K} m_k+\alpha)}{\prod_{k=1}^{K} \Gamma(m_k+\alpha)} \prod_{k=1}^{K} \theta_k^{m_k+\alpha-1} d\theta \\
&= \frac{\Gamma(K\alpha)}{\Gamma(\alpha)^K} \frac{\prod_{k=1}^{K} \Gamma(m_k+\alpha)}{\Gamma(\sum_{k=1}^{K}(m_k+\alpha))}.
\end{aligned} \tag{5}$$

Notice that the part on the right side of the equation on line 4 is a Dirichlet distribution which integrates to 1. Gibbs sampling can be used to approximate the distribution $p(\boldsymbol{Z}|\alpha)$. Let $\boldsymbol{Z_{-(i)}}$ denote the set of hidden labels excluding $z_i$. The probability of $z_i$ can be written as:

$$P(z_i|\boldsymbol{Z_{-(i)}}, \alpha) = \frac{P(z_i, \boldsymbol{Z_{-(i)}}|\alpha)}{P(\boldsymbol{Z_{-(i)}}|\alpha)}. \tag{6}$$

To sample $z_i$, we do not need the exact probability of this equation, but only need to sample from the ratio of the probabilities for each possible value of $z_i$. That is:

$$P(z_i = k|\boldsymbol{Z_{-(i)}}, \alpha) \propto P(z_i = k, \boldsymbol{Z_{-(i)}}|\alpha). \tag{7}$$

Note that the equation on the right takes the form of equation (5) with $z_i$ set equal to $k$ which can be further simplified as follows:

$$\begin{aligned}
P(z_i = k, \boldsymbol{Z_{-(i)}}|\alpha) &= \frac{\Gamma(K\alpha)}{\Gamma(\alpha)^K} \frac{\prod_{k=1}^{K} \Gamma(m_k+\alpha)}{\Gamma(\sum_{k=1}^{K}(m_k+\alpha))} \\
&\propto \prod_{k=1}^{K} \Gamma(m_k+\alpha),
\end{aligned} \tag{8}$$

since the values $\Gamma(K\alpha)$, $\Gamma(\alpha)^K$, and $\Gamma(\sum_{k=1}^{K} m_k + \alpha)$ are all invariant with choice of $z_i$. This equation can be further simplified by using properties of the gamma function and recognizing that the number of items assigned to $k$ will increase by 1 when $z_i$ is set to $k$. We define $m_{k-(i)}$ to be the number of items assigned to $k$ excluding the current $z_i$ that is under investigation.

$$
\begin{aligned}
P(z_i = k, \boldsymbol{Z_{-(i)}}|\alpha) &\propto \prod_{k=1}^{K} \Gamma(m_k + \alpha) \\
&= \Gamma(m_{k-(i)} + \alpha + 1) \prod_{v \neq k} \Gamma(m_{v-(i)} + \alpha) \\
&= (m_{k-(i)} + \alpha)\Gamma(m_{k-(i)} + \alpha) \prod_{v \neq k} \Gamma(m_{v-(i)} + \alpha) \\
&= (m_{k-(i)} + \alpha) \prod_{v=1}^{K} \Gamma(m_{v-(i)} + \alpha) \\
&\propto (m_{k-(i)} + \alpha).
\end{aligned} \tag{9}
$$

Therefore, the probability that item $z_i$ belongs to component $k$ is proportional to the number of items already assigned to component $k$ plus the concentration parameter $\alpha$ (while excluding $z_i$ in the counts). These properties will be used to build collapsed Gibbs samplers for **Celda_C**, **Celda_G**, and **Celda_CG**. We next outline **Celda_C**, the model to cluster cells into populations and estimates the proportions of each population within each sample.

#### 1.2 Generative process

1. For each sample $i \in \{1..S\}$, draw $\theta_i \sim \text{Dir}_K(\alpha)$
2. $\varphi_k \sim \text{Dir}_G(\beta)$ for $k = 1..K$
3. For each cell $j \in \{1..M_i\}$ in sample $i$:
   (a) Draw $z_{i,j} \sim \text{Mult}(\theta_i)$ for $j = 1..M_i$
   (b) For each transcript $t \in \{1..N_{i,j}\}$ in cell $j$ in sample $i$, draw $x_{i,j,t} \sim \text{Mult}(\varphi_{z_{i,j}})$

**Description of parameters:**
$S$ is the number of samples.
$K$ is the number of cell subpopulations.
$G$ is the number of genes.
$M_i$ is the number of cells for sample $i$.
$N_{i,j}$ is the number of transcripts for cell $j$ in sample $i$.
$z_{i,j}$ is the hidden population for cell $j$ in sample $i$ $x_{i,j,t}$ is the $t^{th}$ transcript for cell $j$ in sample $i$

We refer to $\boldsymbol{\theta}$ as the "**Sample Probability (SP)**" matrix as it defines the probability of each cell population in each sample and $\boldsymbol{\varphi}$ as the "**Population Probability (PP)**" matrix as it defines the probability of each gene being observed in each cell population. The complete likelihood function is below followed by the plate diagram (Figure 1):

$$
p(\boldsymbol{X}, \boldsymbol{\theta}, \boldsymbol{Z}, \boldsymbol{\varphi}|\alpha, \beta) = \prod_{i=1}^{S} p(\theta_i|\alpha) \prod_{j=1}^{M_i} p(z_{i,j}|\theta_i) \prod_{k=1}^{K} p(\varphi_k|\beta) \prod_{t=1}^{N_{i,j}} p(x_{i,j,t}|\varphi_{z_{i,j}}). \tag{10}
$$

#### 1.3 Inference using collapsed Gibbs sampling

To build a collapsed Gibbs sampler, $\boldsymbol{\theta}$ and $\boldsymbol{\varphi}$ will be integrated out. As $\boldsymbol{\theta}$ and $\boldsymbol{\varphi}$ are independent, terms dependent on these variables can be grouped and considered separately:

Figure 1: Plate diagram for the cell clustering model Celda_C. Bold circles indicate given prior parameters and shaded circles indicate observed data.

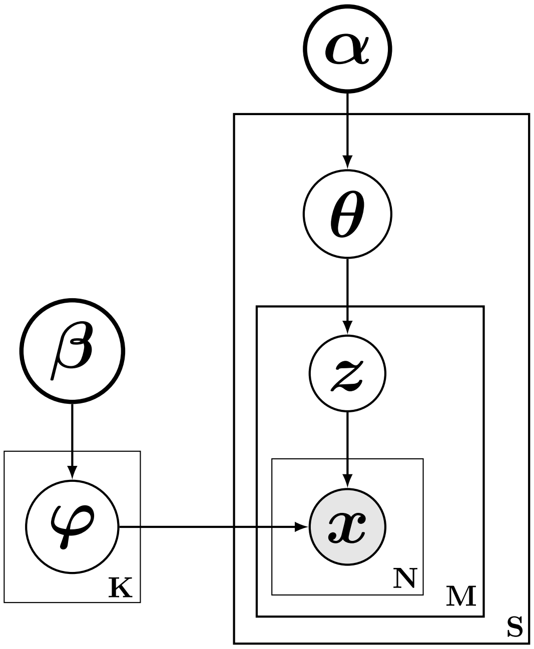


$$
\begin{aligned}
p(\boldsymbol{X}, \boldsymbol{Z}|\alpha, \beta) &= \int_{\boldsymbol{\theta}} \int_{\boldsymbol{\varphi}} p(\boldsymbol{X}, \boldsymbol{\theta}, \boldsymbol{Z}, \boldsymbol{\varphi}|\alpha, \beta) d\boldsymbol{\varphi} d\boldsymbol{\theta} \\
&= \int_{\boldsymbol{\theta}} \int_{\boldsymbol{\varphi}} \prod_{i=1}^{S} p(\theta_i|\alpha) \prod_{j=1}^{M_i} p(z_{i,j}|\theta_i) \prod_{k=1}^{K} p(\varphi_k|\beta) \prod_{t=1}^{N_{i,j}} p(x_{i,j,t}|\varphi_{z_{i,j}}) d\boldsymbol{\varphi} d\boldsymbol{\theta} \\
&= \int_{\boldsymbol{\theta}} \prod_{i=1}^{S} p(\theta_i|\alpha) \prod_{j=1}^{M_i} p(z_{i,j}|\theta_i) d\boldsymbol{\theta} \int_{\boldsymbol{\varphi}} \prod_{k=1}^{K} p(\varphi_k|\beta) \prod_{i=1}^{S} \prod_{j=1}^{M_i} \prod_{t=1}^{N_{i,j}} p(x_{i,j,t}|\varphi_{z_{i,j}}) d\boldsymbol{\varphi} \\
&= \prod_{i=1}^{S} \int_{\theta_i} p(\theta_i|\alpha) \prod_{j=1}^{M_i} p(z_{i,j}|\theta_i) d\theta_i \prod_{k=1}^{K} \int_{\varphi_k} p(\varphi_k|\beta) \prod_{i=1}^{S} \prod_{j \in C_k^i} \prod_{t=1}^{N_{i,j}} p(x_{i,j,t}|\varphi_k) d\varphi_k,
\end{aligned} \tag{11}
$$

where $C_k$ is defined to be the set of cells assigned to subpopulation $k$ and $C_k^i$ is defined to be the set of cells assigned to population $k$ in sample $i$. Note that in the last step, the index for $\varphi$ has been changed from $z_{i,j}$ to $k$ in $P(w_{i,j,t} = y_g|\varphi_k)$ because we have grouped all cells that have the same $k$ rather than ordering them by their index in sample $i$ (i.e. $\prod_{j=1}^{M_i}$). The reording in the last step allows us to focus on the integration for each $\theta_i$ and a $\varphi_k$ separately.

Let $\theta_{i,k}$ be the probability of observing cell population $k$ in sample $i$ and $m_{i,k}$ be the number of cells in sample $i$ that have $z_{i,j} = k$, then the probabilities for cell counts in sample $i$ can be rewritten as:

$$
\prod_{j=1}^{M_i} p(z_{i,j}|\theta_i) = \prod_{k=1}^{K} \theta_{i,k}^{m_{i,k}}. \tag{12}
$$

The procedure outlined by equation (5) can be followed to integrate out each $\theta_i$:

$$
\begin{aligned}
\int_{\theta_i} p(\theta_i|\alpha) \prod_{j=1}^{M_i} p(z_{i,j}|\theta_i) d\theta_i &= \int_{\theta_i} \frac{\Gamma(K\alpha)}{\Gamma(\alpha)^K} \prod_{k=1}^{K} \theta_{i,k}^{\alpha-1} \prod_{k=1}^{K} \theta_{i,k}^{m_{i,k}} d\theta_i \\
&= \frac{\Gamma(K\alpha)}{\Gamma(\alpha)^K} \frac{\prod_{k=1}^{K} \Gamma(m_{i,k}+\alpha)}{\Gamma(\sum_{k=1}^{K}(m_{i,k}+\alpha))}.
\end{aligned} \tag{13}
$$

We can use a similar process to integrate out $\varphi_k$. Let $\varphi_{k,g}$ be the probability of observing gene $g$ in population $k$ and $n_{i,j,g}$ be the number of counts in sample $i$ in cell $j$ for gene $g$, and $n_{(.),(k),g}$ be the sum of all counts across all samples across the subset of cells belonging to population $k$ for gene $g$. The probablity for observed gene counts in population $k$ can be rewritten as:

$$
\prod_{i=1}^{S} \prod_{j \in C_k^i} \prod_{t=1}^{N_{i,j}} p(x_{i,j,t}|\varphi_k) = \prod_{g=1}^{G} \varphi_{k,g}^{n_{(.),(k),g}}. \tag{14}
$$

Now, we can use the procedure outlined in equation (5) to integrate out $\varphi_k$:

$$
\begin{aligned}
\int_{\varphi_k} p(\varphi_k|\beta) \prod_{i=1}^{S} \prod_{j=1}^{M_i} \prod_{t=1}^{N_{i,j}} p(x_{i,j,t}|\varphi_{z_{i,j}}) d\varphi_k &= \int_{\varphi_k} \frac{\Gamma(G\beta)}{\Gamma(\beta)^G} \prod_{g=1}^{G} \varphi_{k,g}^{\beta-1} \prod_{g=1}^{G} \varphi_{k,g}^{n_{(.),(k),g}} d\varphi_k \\
&= \frac{\Gamma(G\beta)}{\Gamma(\beta)^G} \frac{\prod_{g=1}^{G} \Gamma(n_{(.),(k),g}+\beta)}{\Gamma(\sum_{g=1}^{G}(n_{(.),(k),g}+\beta))}.
\end{aligned} \tag{15}
$$

For completeness, the collapsed likelihood with $\boldsymbol{\theta}$ and $\boldsymbol{\varphi}$ integrated out is as follows:

$$
p(\boldsymbol{X}, \boldsymbol{Z}|\alpha, \beta) = \prod_{i=1}^{S} \frac{\Gamma(K\alpha)}{\Gamma(\alpha)^K} \frac{\prod_{k=1}^{K} \Gamma(m_{i,k}+\alpha)}{\Gamma(\sum_{k=1}^{K}(m_{i,k}+\alpha))} \times \prod_{k=1}^{K} \frac{\Gamma(G\beta)}{\Gamma(\beta)^G} \frac{\prod_{g=1}^{G} \Gamma(n_{(.),(k),g}+\beta)}{\Gamma(\sum_{g=1}^{G}(n_{(.),(k),g}+\beta))}. \tag{16}
$$

The distribution $p(\boldsymbol{Z}|\boldsymbol{X}, \alpha, \beta)$ can be approximated with Gibbs sampling. Let $z_{i,j}$ be the hidden subpopulation for cell $j$ in sample $i$, and let $\boldsymbol{Z_{-(i,j)}}$ denote the set of hidden populations for all other cells. We therefore want to derive the following probability:

$$
\begin{aligned}
p(z_{i,j} = k|\boldsymbol{Z_{-(i,j)}}, \boldsymbol{X}, \alpha, \beta) &= \frac{P(z_{i,j} = k, \boldsymbol{Z_{-(i,j)}}, \boldsymbol{X}|\alpha, \beta)}{P(\boldsymbol{Z_{-(i,j)}}, \boldsymbol{X}|\alpha, \beta)} \\
&\propto P(z_{i,j} = k, \boldsymbol{Z_{-(i,j)}}, \boldsymbol{X}|\alpha, \beta).
\end{aligned} \tag{17}
$$

The equation on the right takes the form of the likelihood function with $z_{(i,j)}$ set equal to $k$ which can be simplified according to the procedure outlined in equations (6-9):

$$
\begin{aligned}
P(z_{i,j}=k,\boldsymbol{Z}_{-(\boldsymbol{i},\boldsymbol{j})},\boldsymbol{X}|\alpha,\beta) &= \prod_{i=1}^{S}\frac{\Gamma(K\alpha)}{\Gamma(\alpha)^K}\frac{\prod_{k=1}^{K}\Gamma(m_{i,k}+\alpha)}{\Gamma(\sum_{k=1}^{K}(m_{i,k}+\alpha))}\prod_{k=1}^{K}\frac{\Gamma(G\beta)}{\Gamma(\beta)^G}\frac{\prod_{g=1}^{G}\Gamma(n_{(.),(k),g}+\beta)}{\Gamma(\sum_{g=1}^{G}(n_{(.),(k),g}+\beta))} \\
&= \left[\frac{\Gamma(K\alpha)}{\Gamma(\alpha)^K}\right]^S\prod_{i=1}^{S}\frac{\prod_{k=1}^{K}\Gamma(m_{i,k}+\alpha)}{\Gamma(\sum_{k=1}^{K}(m_{i,k}+\alpha))}\left[\frac{\Gamma(G\beta)}{\Gamma(\beta)^G}\right]^K\prod_{k=1}^{K}\frac{\prod_{g=1}^{G}\Gamma(n_{(.),(k),g}+\beta)}{\Gamma(\sum_{g=1}^{G}(n_{(.),(k),g}+\beta))} \\
&\propto \prod_{i=1}^{S}\frac{\prod_{k=1}^{K}\Gamma(m_{i,k}+\alpha)}{\Gamma(\sum_{k=1}^{K}(m_{i,k}+\alpha))}\prod_{k=1}^{K}\frac{\prod_{g=1}^{G}\Gamma(n_{(.),(k),g}+\beta)}{\Gamma(\sum_{g=1}^{G}(n_{(.),(k),g}+\beta))} \\
&\propto \prod_{k=1}^{K}\Gamma(m_{i,k}+\alpha)\prod_{k=1}^{K}\frac{\prod_{g=1}^{G}\Gamma(n_{(.),(k),g}+\beta)}{\Gamma(\sum_{g=1}^{G}(n_{(.),(k),g}+\beta))} \\
&\propto (m_{i,k-(i,j)}+\alpha)\prod_{k=1}^{K}\frac{\prod_{g=1}^{G}\Gamma(n_{(.),(k),g}+\beta)}{\Gamma(\sum_{g=1}^{G}(n_{(.),(k),g}+\beta))},
\end{aligned}
\tag{18}
$$

where $m_{i,k}$ is the number of cells assigned to population $k$ in sample $i$ and $m_{i,k-(i,j)}$ is the number of cells assigned to population $k$ in sample $i$ excluding the cell $j$ in sample $i$. The left side of the equation could further simplified in line 3 due to the fact that the label is only changing for cell $j$ in sample $i$ and the number of cells in each population for all other samples will be stationary. Therefore, the part of the equation concerning the counts for all samples other than sample $i$ will be invariant with respect to $z_{i,j}$. For clarity:

$$
\begin{aligned}
\prod_{i=1}^{S}\frac{\prod_{k=1}^{K}\Gamma(m_{i,k}+\alpha)}{\Gamma(\sum_{k=1}^{K}(m_{i,k}+\alpha))} &= \frac{\prod_{k=1}^{K}\Gamma(m_{i,k}+\alpha)}{\Gamma(\sum_{k=1}^{K}(m_{i,k}+\alpha))}\prod_{v\neq i}\frac{\prod_{k=1}^{K}\Gamma(m_{v,k}+\alpha)}{\Gamma(\sum_{k=1}^{K}(m_{v,k}+\alpha))} \\
&\propto \prod_{k=1}^{K}\Gamma(m_{i,k}+\alpha),
\end{aligned}
\tag{19}
$$

where $v$ is the set of sample indices that are not equal to the current sample $i$. After completing Gibbs sample and identifying the $\boldsymbol{Z}$ with the highest probability, point estimates for the Dirichlet distribution probabilities can derived as described in the next section.

#### 1.4 Inference using point estimates

Given sample of $\boldsymbol{Z}$, we can derive point estimates for the Dirichlet distributions. For $\boldsymbol{\theta}$, we have:

$$
\hat{\theta}_{i,k} = \frac{m_{i,k}+\alpha}{M_i+K\alpha}, \tag{20}
$$

where $m_{i,k}$ is the number of cells assigned to population $k$ in sample $i$ and $M_i$ is the total number of cells in sample $i$. For $\boldsymbol{\varphi}$, we have:

$$
\hat{\varphi}_{k,g} = \frac{n_{(\cdot),(k),g}+\beta}{n_{(\cdot),(k),(\cdot)}+G\beta}, \tag{21}
$$

where $n_{(\cdot),(k),g}$ is the sum of counts across all samples and cells belonging to subpopulaiton $k$ for gene $g$. $n_{(\cdot),(k),(\cdot)}$ is defined as the sum of counts across all samples and across all cells in subpopulation $k$ for all genes.

Given $\hat{\boldsymbol{\theta}}$ and $\hat{\boldsymbol{\varphi}}$, we can identify the most likely cluster label for each $z_{i,j}$:

$$\begin{aligned}\hat{z}_{i,j} &= \text{argmax}_k \left\{ p(z_{i,j} = k | \boldsymbol{X}_{i,j}, \hat{\boldsymbol{\theta}}, \hat{\boldsymbol{\varphi}}) \right\} \\ &= \text{argmax}_k \left\{ \hat{\theta}_{i,k} \prod_{g=1}^{g} \hat{\varphi}_{k,g}^{n_{i,j,g}} \right\},\end{aligned} \tag{22}$$

where $\boldsymbol{X}_{i,j}$ is the counts for cell $j$ in sample $i$. In this inference procedure, we alternate between estimating the Dirichlet distribution probabilities ($\hat{\boldsymbol{\theta}}$ and $\hat{\boldsymbol{\varphi}}$) and the cell labels ($\hat{z}_{i,j}$). Since each cell is fully assigned to a subpopulation, this procedure is not truly expectation-maximization (EM). It more closely resembles the procedure used in K-means clustering, which is sometimes refered to as "hard" EM.

##### 1.5 Perplexity

Perplexity is a measure directly related to the probability of observing cell counts given the estimated model parameters and can be used in cross validation or subsampling to help in the choice of $K$. Perplexity is defined as:

$$\text{Perplexity(x)} = \exp\left( -\frac{\log(p(x))}{\sum_{i=1}^{S} \sum_{j=1}^{M_i} N_{i,j}} \right), \tag{23}$$

where $N_{i,j}$ is the number of counts within each cell $j$ in sample $i$. $\log(p(x))$ is defined as:

$$\log(p(x)) = \sum_{i=1}^{S} \sum_{j=1}^{M_j} \log \left[ \sum_{k=1}^{K} \theta_{i,k} \prod_{g=1}^{G} \varphi_{k,g}^{n_{i,j,g}} \right], \tag{24}$$

where $\theta_{i,k}$ is the probabiity of a cell belonging to a cell population $k$ in sample $i$, $\varphi_{k,g}$ is the probability of gene $g$ in cell population $k$, $n_{j,g}$ is the count of gene $g$ in cell $j$ in sample $i$.

### 2 Celda_G: Clustering genes into transcriptional modules across cells

#### 2.1 Background

The goal of text mining models such as Latent Dirichlet Allocation (LDA) is to identify hidden components called "topics" across a set of documents and estimate the degree to which each topic is present in each document [3]. Each topic is a distribution over words in the vocabulary and represents the degree to which the frequency of word counts co-occur with each other across the documents. Furthuremore, each document is treated as a distribution over the collection of topics and with each document having a different combination of topics. The goal of the **Celda_G** model is to cluster genes into "transcriptional modules" and estimate the degree to which each module is present in each cell. The fundemental biological principle being leveraged is that genes which are under control of the same transcriptional programs (i.e. transcription factors and epigenetic regulators) will co-vary in expression across cells.

The goal of many gene expression clustering algorithms is to group genes into distinct, non-overlapping sets of genes (i.e. hard-clustering of genes). The rationale for this type of clustering is that genes with highly similar expression patterns will be involved in the same biological processes. In LDA and the majority of topic models, each word has a non-zero probability in each topic and therefore every topic can emit every word. As such, topics can be viewed as a mixture of words, a form of "soft-clustering". In the context of gene expression, this can lead to difficulties in interpretation as all genes will belong to all transcriptional modules. For example it would be difficult to interpret a transcriptional module that had a non-zero probability for ciliary- and adipose-related genes. One could apply various post-hoc analyses to the results of LDA to find which types of genes have relatively higher probabilities in each transcriptional module. However, this requires the choice of which additional heuristic to use along with its various thresholds. We sought to streamline the inference by incorporating the hard-clustering directly into the model and assigning each gene to a unique transcriptional module.

The sparse Topic Model (sparseTM) is an extension to LDA that has the capability to turn words completely "off" in different topics [4]. The goal of the sparseTM is to decouple sparsity and smoothness in the topic distributions. While words still have the capability to be emitted by more than one topic, each word will not necessarily be emitted by all topics. Here, we leverage this technique to turn off genes in all transcriptional modules except one, thus performing "hard-clustering". For most topic models, the topics are drawn from an exchangeable Dirichlet in which the components of the vector parameter are equal to the same scalar. Assume that $G$ is the number of genes. The exchangeable Dirichlet for a transcriptional module, $\psi$, can be described as:

$$\psi \sim \text{Dir}_G(\delta \mathbf{1}), \tag{25}$$

where $\psi$ is drawn from a Dirichlet parameterized by the scalar $\delta$ multiplied by a $G$-length vector of 1's. In a Dirichlet distribution, if a parameter is set to zero instead of a positive scalar, the probability of that component in the resulting distribution will also be zero. In the Celda_G model, we use different indicator vectors for each transcriptional module to turn genes "on" or "off" in each module:

$$\psi_l \sim \text{Dir}_G(\delta \boldsymbol{Y}_l). \tag{26}$$

Here $\boldsymbol{Y}_l$ is a $G$-length vector of 0's and 1's for transcriptional module $\psi_l$ used to set each parameter of the Dirichlet to 0 or $\delta$. In contrast to sparseTM, the $\boldsymbol{Y}_l$ vectors are controlled such that a gene is only turned on in a single module which will result in distinct, non-overlapping clusters of genes. The entire generative process is outlined in the next section.

#### 2.2 Generative process

1. Draw $\eta \sim \text{Dir}_L(\gamma)$
2. For each gene $g \in \{1..G\}$, draw $y_g \sim \text{Mult}(\eta)$
3. For each transcriptional module distribution $l \in \{1..L\}$ :
   (a) Define $Y_l = [y_g = l]_{g=1}^{G}$

(b) Draw $\psi_l \sim \text{ Dir}_G(\delta Y_l)$

4. For each cell $j \in \{1..M\}$:
   (a) Draw transcriptional module proportions $\varphi_j \sim \text{ Dir}_L(\beta)$
   (b) For the $t$-th transcript in cell $j$, $t \in \{1..N_j\}$:
      i. Draw $w_{j,t} \sim \text{ Mult}(\varphi_j)$
      ii. Draw $x_{j,t} \sim \text{ Mult}(\psi_{w_{j,t}})$

**Description of parameters:**
$L$ is the number of transcriptional modules.
$G$ is the number of genes.
$y_g$ is the hidden module for gene $g$
$M$ is the number of cells.
$N_j$ is the number of transcripts for cell $j$.
$w_{j,t}$ is the hidden module for transcript $x_{j,t}$
$x_{j,t}$ is the $t^{th}$ transcript for cell $j$

In this model, $\eta$ is a Dirichlet with length equal to the total number of transcriptional modules specified by $L$. $y_g$ is a single categorical draw from $\eta$ for gene $g$ and will return a value between 1 and $L$. "[ ]" refers to a Boolean operator and returns 1 when the expression within the bracket is true and 0 otherwise. We use this operator in step 3a to denote that the component in $Y_l$ corresponding to gene $g$ will be set to 1 if $y_g = l$ and 0 otherwise. $Y_l$ will then be used as an indicator variable in step 3b to control the genes turned on in transcriptional module $l$. The combination of these variables is used to control the assignment of each gene to a single transcriptional module.

We also note that this model has two levels of hidden variables, one for the overall gene, $y_g$, and one for each individual transcript $t$ in each cell $j$, $w_{j,t}$. Another interesting property of this framework is that the posterior will have a probability of 0 anywhere the hidden variable for an individual count of a gene does not equal the hidden variable for the overall gene. This property will be utilized to marginalize out the set of $W$ as decribed in the inference section.

We also implement a similar nuance introduced in the sparseTM by Wang and Blei et al (2009). When a transcriptional module does not have any genes assigned to it, the likelihood is undefined. To overcome this problem, an additional "auxiliary" gene is introduced as the "$G + 1$"-th term in the matrix and assigned to the module with no genes. Otherwise, the indicator variable for the auxiliary gene will be set to 0 in all modules. Note that the auxiliary genes do not have any observed counts in the dataset. We allow for multiple auxiliary genes to be instantiated if multiple transcriptional modules do not have any genes assigned to them.

We refer to $\boldsymbol{\varphi}$ as the "**Module Probability (MP)**" matrix as it defines the probablity of each transcriptional module within each cell and $\boldsymbol{\psi}$ as the "**Gene Probability (GP)**" matrix as it defines the probability of each gene within each transcriptional module. The complete likelihood function is below followed by the plate diagram (Figure 2):

$$P(\eta, \boldsymbol{\varphi}, \boldsymbol{\psi}, \boldsymbol{Y}, \boldsymbol{W}, \boldsymbol{X}|\beta, \delta, \gamma) = P(\eta|\gamma) \prod_{g=1}^{G} P(y_g|\eta) \prod_{j=1}^{M} P(\varphi_j|\beta) \prod_{l=1}^{L} P(\psi_l|\delta, Y_l) \prod_{t=1}^{N_j} P(w_{j,t}|\varphi_j) P(x_{j,t}|\psi_{w_{j,t}}). \tag{27}$$

#### 2.3 Inference using collapsed Gibbs sampling

To build a collapsed Gibbs sampler, we will integrate out $\eta$, $\boldsymbol{\varphi}$, $\boldsymbol{\psi}$, and $\boldsymbol{W}$:

Figure 2: Plate diagram for gene clustering model, Celda_G. Bold circles indicate given prior parameters and shaded circles indicate observed data.

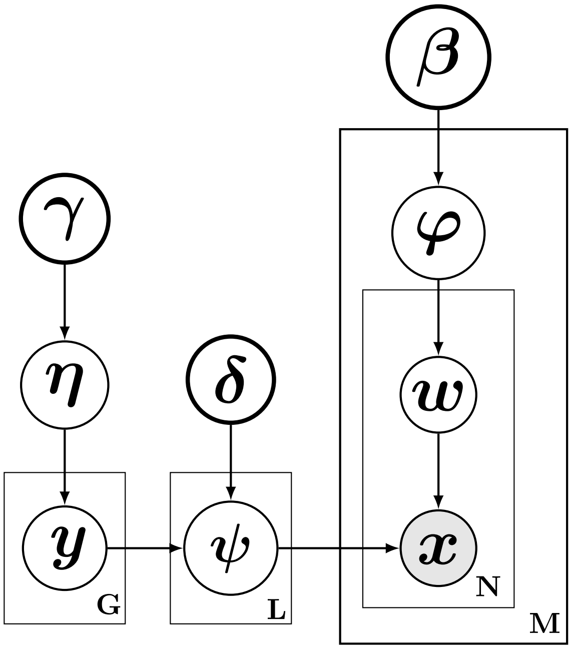


$$
\begin{aligned}
P(\boldsymbol{Y}, \boldsymbol{X}|\beta, \delta, \gamma) = \int_{\boldsymbol{\eta}} \int_{\boldsymbol{\varphi}} \int_{\boldsymbol{\psi}} P(\eta|\gamma) \prod_{g=1}^{G} P(y_g|\eta) \prod_{j=1}^{M} P(\varphi_j|\beta) \prod_{l=1}^{L} P(\psi_l|\delta, Y_l) \\
\times \prod_{t=1}^{N_j} \left( \sum_{v=1}^{L} P(w_{j,t} = v|\varphi_j) P(x_{j,t} = g|\psi_v) \right) d\boldsymbol{\psi} d\boldsymbol{\varphi} d\eta.
\end{aligned} \tag{28}
$$

We will first simplify the marginalization over the hidden state for each count, $w_{j,t}$. This sum can be subdivived into two components. The first component corresponds to part of the summation where the hidden state assignment for an individual count within a gene (denoted by $w_{j,t}$) is the same as the overall hidden gene label (denoted by $y_g$). The second part corresponds to summation over the remaining components where the hidden state assignment for the count is not equal to the overall gene label $y_g$:

$$
\begin{aligned}
\sum_{v=1}^{L} P(w_{j,t} = v|\varphi_j) P(x_{j,t} = g|\psi_v) = P(w_{j,t} = y_g|\varphi_j) P(x_{j,t} = g|\psi_{y_g}) \\
+ \sum_{v' \neq y_g} P(w_{j,t} = v'|\varphi_j) P(x_{j,t} = g|\psi_{v'}).
\end{aligned} \tag{29}
$$

In LDA, all of the components in this marginalization would be nonzero. However, in this model, genes have been assigned to a single transcriptional module based on the set of $\boldsymbol{Y}$ indicator variables. That is, $P(x_{j,t} = g|\psi_v) = 0$ when $v$ is not equal to $y_g$ for gene $g$. In other words, since each gene can only be assigned to a single transcriptional module, according to $y_g$, the probability of a count being assigned to that gene in another transcriptional module (i.e. when $w_{j,t} \neq y_g$) is zero. In fact, the only nonzero component in the sum will occur when $w_{j,t} = y_g$. The combination of the $\boldsymbol{Y}$ indicator variables and the marginalization of the $\boldsymbol{W}$ allows us to simultaneously estimate the same hidden state for all counts in a gene rather than sampling different states for each individual count and thus provides the "hard clustering" behavior on the genes. For clarity, the sum is simplified to:

$$\sum_{v=1}^{L} P(w_{j,t} = v|\varphi_j)P(x_{j,t} = g|\psi_v) = P(w_{j,t} = y_g|\varphi_j)P(x_{j,t} = g|\psi_{y_g}). \tag{30}$$

And the overall likelihood will "collapse" back down to a product:

$$\begin{aligned} &P(\boldsymbol{Y}, \boldsymbol{X}|\beta, \delta, \gamma) = \\ &\int_{\boldsymbol{\eta}}\int_{\boldsymbol{\varphi}}\int_{\boldsymbol{\psi}} P(\eta|\gamma)\prod_{g=1}^{G} P(y_g|\eta)\prod_{j=1}^{M} P(\varphi_j|\beta)\prod_{l=1}^{L} P(\psi_l|\delta, Y_l)\prod_{t=1}^{N_j} P(w_{j,t} = y_g|\varphi_j)P(x_{j,t} = g|\psi_{y_g})d\boldsymbol{\psi} d\boldsymbol{\varphi} d\eta. \end{aligned} \tag{31}$$

We can also integrate out the probabilities from the Dirichlet-multinomial distributions. First, we group terms related to each integration:

$$\begin{aligned} P(\boldsymbol{Y}, \boldsymbol{X}|\beta, \delta, \gamma) = &\int_{\eta} P(\eta|\gamma)\prod_{g=1}^{G} P(y_g|\eta)d\eta \\ &\times \int_{\boldsymbol{\varphi}}\prod_{j=1}^{M} P(\varphi_j|\beta)\prod_{t=1}^{N_j} P(w_{j,t} = y_g|\varphi_j)d\boldsymbol{\varphi} \\ &\times \int_{\boldsymbol{\psi}}\prod_{l=1}^{L} P(\psi_l|\delta, Y_l)\prod_{j=1}^{M}\prod_{t=1}^{N_j} P(x_{j,t} = g|\psi_{y_g})d\boldsymbol{\psi}. \end{aligned} \tag{32}$$

If we use the notation $V_l$ to denote the subset of genes that are assigned to module $l$ and $|V_l|$ to denote the number of genes assigned to module $l$, we can re-write the series of multinomial probabilties related to $\eta$:

$$\prod_{g=1}^{G} p(y_g|\eta) = \prod_{l=1}^{L} \eta_l^{|V_l|}, \tag{33}$$

where $\eta_l$ is the probability of a gene being assigned to module $l$. Now $\eta$ can be integrated out according to equation (5):

$$\begin{aligned} \int_{\eta} P(\eta|\gamma)\prod_{g=1}^{G} P(y_g|\eta)d\eta &= \int_{\eta} \frac{\Gamma(L\gamma)}{\Gamma(\gamma)^L}\prod_{l=1}^{L}\eta_l^{\gamma-1}\prod_{l=1}^{L}\eta_l^{|V_l|}d\eta \\ &= \frac{\Gamma(L\gamma)}{\Gamma(\gamma)^L}\frac{\prod_{l=1}^{L}\Gamma(|V_l| + \gamma)}{\Gamma(\sum_{l=1}^{L}(|V_l| + \gamma))}. \end{aligned} \tag{34}$$

Next, we focus on $\boldsymbol{\varphi}$. Let $n_{j,g}$ be the number of counts for gene $g$ in cell $j$ and let $(\cdot)$ be used to indicate a sum across all elements in a dimension. Also, let $n_{j,(V_l)}$ be the sum of counts from the set of genes assigned to module $l$ in cell $j$. We can re-write the multinomial probabilities for a single cell $j$ as:

$$\prod_{t=1}^{N_j} P(w_{j,t} = y_g|\varphi_j) = \prod_{l=1}^{L} \varphi_{j,l}^{n_{j,(V_l)}}, \tag{35}$$

where $\varphi_{j,l}$ is the probability of a module $l$ in cell $j$. Now terms can be grouped and each $\varphi_l$ can be integrated out according to equation (5):

$$\begin{aligned}
\int_{\boldsymbol{\varphi}} \prod_{j=1}^{M} P(\varphi_j|\beta) \prod_{t=1}^{N_j} P(w_{j,t} = y_g|\varphi_j) d\boldsymbol{\varphi} &= \int_{\boldsymbol{\varphi}} \prod_{j=1}^{M} \frac{\Gamma(L\beta)}{\Gamma(\beta)^L} \prod_{l=1}^{L} \varphi_{j,l}^{\beta-1} \prod_{l=1}^{L} \varphi_{j,l}^{n_{j,(V_l)}} d\boldsymbol{\varphi} \\
&= \prod_{j=1}^{M} \int_{\varphi_j} \frac{\Gamma(L\beta)}{\Gamma(\beta)^L} \prod_{l=1}^{L} \varphi_{j,l}^{\beta-1} \prod_{l=1}^{L} \varphi_{j,l}^{n_{j,(V_l)}} d\varphi_j \\
&= \prod_{j=1}^{M} \frac{\Gamma(L\beta)}{\Gamma(\beta)^L} \frac{\prod_{l=1}^{L} \Gamma(n_{j,(V_l)} + \beta)}{\Gamma(\sum_{l=1}^{L}(n_{j,(V_l)} + \beta))}.
\end{aligned} \tag{36}$$

Let, $n_{(\cdot),g}$ represent the sum of counts for gene $g$ across all cells. The multinomial probabilities related to $\psi_l$ can be re-written as:

$$\prod_{j=1}^{M} \prod_{t=1}^{N_j} P(x_{j,t} = g|\psi_{y_g}) = \prod_{l=1}^{L} \prod_{v \in V_l} \psi_{l,v}^{n_{(\cdot),v}}. \tag{37}$$

We use the notation $\sum_{v \in V_l}$ or $\prod_{v \in V_l}$ to denote the sum or product over genes assigned to module $l$, respectively. That is, the sum or product over the active, nonzero components of $\psi_l$.

$$\begin{aligned}
\int_{\boldsymbol{\psi}} P(\psi_l|\delta, Y_l) \prod_{j=1}^{M} \prod_{t=1}^{N_j} P(x_{j,t} = g|\psi_{y_g}) d\boldsymbol{\psi} &= \int_{\boldsymbol{\psi}} \prod_{l=1}^{L} \left( \frac{\Gamma(|V_l|\delta)}{\Gamma(\delta)^{|V_l|}} \prod_{v \in V_l} \psi_{l,v}^{\delta-1} \right) \prod_{l=1}^{L} \left( \prod_{v \in V_l} \psi_{l,v}^{n_{(\cdot),v}} \right) d\boldsymbol{\psi} \\
&= \prod_{l=1}^{L} \int_{\psi_l} \frac{\Gamma(|V_l|\delta)}{\Gamma(\delta)^{|V_l|}} \prod_{v \in V_l} \psi_{l,v}^{\delta-1} \prod_{v \in V_l} \psi_{l,v}^{n_{(\cdot),v}} d\psi_l \\
&= \prod_{l=1}^{L} \frac{\Gamma(|V_l|\delta)}{\Gamma(\delta)^{|V_l|}} \frac{\prod_{v \in V_l} \Gamma(n_{(\cdot),v} + \delta)}{\Gamma(\sum_{v \in V_l}(n_{(\cdot),v} + \delta))}.
\end{aligned} \tag{38}$$

In summary, the complete collapsed likelihood is as follows:

$$\begin{aligned}
P(\boldsymbol{Y}, \boldsymbol{X}|\beta, \delta, \gamma) = &\frac{\Gamma(L\gamma)}{\Gamma(\gamma)^L} \times \frac{\prod_{l=1}^{L} \Gamma(|V_l| + \gamma)}{\Gamma(\sum_{l=1}^{L}(|V_l| + \gamma))} \\
&\times \prod_{j=1}^{M} \frac{\Gamma(L\beta)}{\Gamma(\beta)^L} \times \frac{\prod_{l=1}^{L} \Gamma(n_{j,(V_l)} + \beta)}{\Gamma(\sum_{l=1}^{L}(n_{j,(V_l)} + \beta))} \\
&\times \prod_{l=1}^{L} \frac{\Gamma(|V_l|\delta)}{\Gamma(\delta)^{|V_l|}} \times \frac{\prod_{v \in V_l} \Gamma(n_{(\cdot),v} + \delta)}{\Gamma(\sum_{v \in V_l}(n_{(\cdot),v} + \delta))}.
\end{aligned} \tag{39}$$

The distribution $p(\boldsymbol{Y}|\boldsymbol{X}, \beta, \delta, \gamma)$ can be approximated with Gibbs sampling. Let $y_g$ be the hidden state for gene $j$ and let $\boldsymbol{Y}_{-(g)}$ denote the set of hidden modules for all other genes. We therefore want to derive the following probability:

$$
\begin{aligned}
P(y_g = l|\boldsymbol{Y}_{-(g)}, \boldsymbol{X}, \beta, \delta, \gamma) &= \frac{P(y_g = l, \boldsymbol{Y}_{-(g)}, \boldsymbol{X}|\beta, \delta, \gamma)}{P(\boldsymbol{Y}_{-(g)}, \boldsymbol{X}|\beta, \delta, \gamma)} \\
&\propto P(y_g = l, \boldsymbol{Y}_{-(g)}, \boldsymbol{X}|\beta, \delta, \gamma),
\end{aligned} \tag{40}
$$

which is equal to the likelihood equation listed above. The likelihood equation can be simplified by removing elements that are invariant with different choices of $y_g$. For the first component, we have

$$
\frac{\Gamma(L\gamma)}{\Gamma(\gamma)^L} \times \frac{\prod_{l=1}^{L} \Gamma(|V_l| + \gamma)}{\Gamma(\sum_{l=1}^{L}(|V_l| + \gamma))} \propto \frac{\prod_{l=1}^{L} \Gamma(|V_l| + \gamma)}{\Gamma(\sum_{l=1}^{L}(|V_l| + \gamma))}. \tag{41}
$$

For the second component, we have:

$$
\begin{aligned}
\prod_{j=1}^{M} \frac{\Gamma(L\beta)}{\Gamma(\beta)^L} \times \frac{\prod_{l=1}^{L} \Gamma(n_{j,(V_l)} + \beta)}{\Gamma(\sum_{l=1}^{L}(n_{j,(V_l)} + \beta))} &= \left[\frac{\Gamma(L\beta)}{\Gamma(\beta)^L}\right]^M \times \frac{\prod_{j=1}^{M}\prod_{l=1}^{L} \Gamma(n_{j,(V_l)} + \beta)}{\prod_{j=1}^{M} \Gamma(\sum_{l=1}^{L}(n_{j,(V_l)} + \beta))} \\
&\propto \prod_{j=1}^{M}\prod_{l=1}^{L} \Gamma(n_{j,(V_l)} + \beta).
\end{aligned} \tag{42}
$$

Note that $\sum_{l=1}^{L} n_{j,(V_l)} + \beta = N_j + L\beta$, where $N_j$ is the total number of counts in cell $j$. This quantity is invariant with respect to choice of $y_g$ and thus can be dropped. For the final component, we have:

$$
\prod_{l=1}^{L} \frac{\Gamma(|V_l|\delta)}{\Gamma(\delta)^{|V_l|}} \times \frac{\prod_{v \in V_l} \Gamma(n_{(\cdot),v} + \delta)}{\Gamma(\sum_{v \in V_l}(n_{(\cdot),v} + \delta))} \propto \prod_{l=1}^{L} \frac{\Gamma(|V_l|\delta)}{\Gamma(\delta)^{|V_l|}} \frac{1}{\Gamma(\sum_{v \in V_l}(n_{(\cdot),v} + \delta))}. \tag{43}
$$

The full Gibbs sampling equation is as follows:

$$
P(y_g = l|\boldsymbol{X}, \boldsymbol{Y}_{-(g)}, \beta, \delta, \gamma) \propto \frac{\prod_{l=1}^{L} \Gamma(|V_l| + \gamma)}{\Gamma(\sum_{l=1}^{L}(|V_l| + \gamma))} \times \prod_{j=1}^{M}\prod_{l=1}^{L} \Gamma(n_{j,(V_l)} + \beta) \times \prod_{l=1}^{L} \frac{\Gamma(|V_l|\delta)}{\Gamma(\delta)^{|V_l|}} \frac{1}{\Gamma(\sum_{v \in V_l}(n_{(\cdot),v} + \delta))}. \tag{44}
$$

This equation could be further simplified when no auxillary genes are instantiated. For example, the quantity $\prod_{l=1}^{L} \Gamma(\delta)^{|V_l|} = \Gamma(\delta)^{\sum_l |V_l|} = \Gamma(\delta)^G$ is invariant to the choice of $y_g$ when no auxillary genes are instantiated. However, when an auxillary gene is activated in a module without any real genes, the total number of genes $G$ increases by one. Therefore, we did not simplify components that depend on the total number of genes further.

The second and third terms of above equation can be further simplified to speed up computing time as follows, given that gene $g$ is currently in module $l$:

$$
\begin{aligned}
\prod_{j=1}^{M}\prod_{l=1}^{L}\Gamma(n_{j,(V_l)}+\beta) &= \prod_{j=1}^{M}\left[\prod_{l=1}^{L}\Gamma(n_{j,(V_l)}+\beta)\right]\\
&= \prod_{j=1}^{M}\left[\Gamma(n_{j,(V_l)}+\beta)\times\prod_{l':l'\neq l}\Gamma(n_{j,(V_{l'})}+\beta)\right]\\
&= \prod_{j=1}^{M}\left[\Gamma(n_{j,(V_l)}+\beta)\times\prod_{l':l'\neq l}\Gamma(n^{-(g)}_{j,(V_{l'})}+\beta)\right]\\
&= \prod_{j=1}^{M}\left[\frac{\Gamma(n_{j,(V_l)}+\beta)}{\Gamma(n^{-(g)}_{j,(V_l)}+\beta)}\times\prod_{l'=1}^{L}\Gamma(n^{-(g)}_{j,(V_{l'})}+\beta)\right]\\
&\propto \prod_{j=1}^{M}\left[\frac{\Gamma(n_{j,(V_l)}+\beta)}{\Gamma(n^{-(g)}_{j,(V_l)}+\beta)}\right]
\end{aligned}
\tag{45}
$$

$$
\begin{aligned}
\prod_{l=1}^{L}\frac{\Gamma(|V_l|\delta)}{\Gamma(\delta)^{|V_l|}}\frac{1}{\Gamma(\sum_{v\in V_l}(n_{(\cdot),v}+\delta))} &= \prod_{l=1}^{L}\frac{\Gamma(|V_l|\delta)}{\Gamma(\delta)^{|V_l|}}\times\prod_{l=1}^{L}\frac{1}{\Gamma(\sum_{v\in V_l}(n_{(\cdot),v}+\delta))}\\
&= \left[\prod_{l=1}^{L}\frac{\Gamma(|V_l|\delta)}{\Gamma(\delta)^{|V_l|}}\right]\times\left[\frac{1}{\Gamma(\sum_{v\in V_l}(n_{(\cdot),v}+\delta))}\prod_{l':l'\neq l}\frac{1}{\Gamma(\sum_{v\in V_{l'}}(n_{(\cdot),v}+\delta))}\right]\\
&= \left[\prod_{l=1}^{L}\frac{\Gamma(|V_l|\delta)}{\Gamma(\delta)^{|V_l|}}\right]\times\left[\frac{1}{\Gamma(\sum_{v\in V_l}(n_{(\cdot),v}+\delta))}\prod_{l':l'\neq l}\frac{1}{\Gamma(\sum_{v\in V_{l'}}(n^{-(g)}_{(\cdot),v}+\delta))}\right]\\
&= \left[\prod_{l=1}^{L}\frac{\Gamma(|V_l|\delta)}{\Gamma(\delta)^{|V_l|}}\right]\times\left[\frac{\Gamma(\sum_{v\in V_l}(n^{-(g)}_{(\cdot),v}+\delta))}{\Gamma(\sum_{v\in V_l}(n_{(\cdot),v}+\delta))}\prod_{l'=1}^{L}\frac{1}{\Gamma(\sum_{v\in V_{l'}}(n^{-(g)}_{(\cdot),v}+\delta))}\right]\\
&\propto \left[\prod_{l=1}^{L}\frac{\Gamma(|V_l|\delta)}{\Gamma(\delta)^{|V_l|}}\right]\times\left[\frac{\Gamma(\sum_{v\in V_l}(n^{-(g)}_{(\cdot),v}+\delta))}{\Gamma(\sum_{v\in V_l}(n_{(\cdot),v}+\delta))}\right],
\end{aligned}
$$

where $n^{-(g)}$ is the total number of transcripts leaving out those from gene $g$. Specifically, for example, $n^{-(g)}_{j,(V_l)}$ is the total number of transcripts in cell $j$ of all the genes in module $l$ leaving out those from gene $g$. Note that when gene $g$ is currently not in module $l'$, $n^{-(g)}_{j,(V'_l)}$ is the same as $n_{j,(V'_l)}$.

Hence the full Gibbs sampling equation is simplified as:

$$
P(y_g=l|\boldsymbol{X},\boldsymbol{Y}_{-(g)},\beta,\delta,\gamma)\propto\frac{\prod_{l=1}^{L}\Gamma(|V_l|+\gamma)}{\Gamma(\sum_{l=1}^{L}(|V_l|+\gamma))}\times\prod_{j=1}^{M}\left[\frac{\Gamma(n_{j,(V_l)}+\beta)}{\Gamma(n^{-(g)}_{j,(V_l)}+\beta)}\right]\times\left[\prod_{l=1}^{L}\frac{\Gamma(|V_l|\delta)}{\Gamma(\delta)^{|V_l|}}\right]\times\frac{\Gamma(\sum_{v\in V_l}(n^{-(g)}_{(\cdot),v}+\delta))}{\Gamma(\sum_{v\in V_l}(n_{(\cdot),v}+\delta))}.
\tag{46}
$$

#### 2.4 Posterior point estimates for Dirichlet distributions

With given sample of $\boldsymbol{Y}$, we can derive point estimates for the Dirichlet distributions. For $\boldsymbol{\varphi}$, we have:

$$
\hat{\varphi}_{j,l}=\frac{n_{j,(V_l)}+\beta}{N_j+L\beta},
\tag{47}
$$

where $n_{j,(V_l)}$ is the sum of counts across all genes belonging to module $l$ for cell $j$ and $N_j$ is the total number of counts for cell $j$. For $\boldsymbol{\psi}$, we have:

$$\hat{\psi}_{l,g} = \frac{n_{(\cdot),g} + \delta}{n_{(\cdot),(V_l)} + |V_l|\delta}, \tag{48}$$

if $y_g = l$ (i.e. if gene $g$ is assigned to transcriptional module $l$). If $y_g \neq l$, then the posterior probability is 0. $n_{(\cdot),g}$ is the sum of counts for gene $g$ across all cells and $n_{(\cdot),(V_l)}$ is the sum of counts across all genes belonging to module $l$ for all cells.

##### 2.5 Perplexity

Perplexity was previously defined in equation (23). For the Celda_G model, we define $\log(p(x))$ to be:

$$\log(p(x)) = \sum_{j=1}^{M_j} \sum_{g=1}^{G} n_{j,g} \log \left[ \sum_{l=1}^{L} \varphi_{j,l} \psi_{l,g} \right], \tag{49}$$

where $\varphi_{j,l}$ is the probabiity transcriptional module $l$ in cell $j$, $\psi_{l,g}$ is the probability of gene $g$ in transcriptional module $l$, $n_{j,g}$ is the number of counts in gene $g$ for cell $j$, and $\eta_l$ is the probability of a gene being assigned to transcriptional module $l$. Note that the only non-zero $\psi_{l,g}$ will be where $y_g = l$ for gene $g$ and thus the sum of the equation can be simplified to:

$$\log(p(x)) = \sum_{j=1}^{M_j} \sum_{g=1}^{G} n_{j,g} \log(\varphi_{j,y_g} \psi_{y_g,g}). \tag{50}$$

### 3 Celda_CG: Simultaneous clustering of genes into transcriptional modules and cells into subpopulation

#### 3.1 Background

**Celda_CG** combines principles from both **Celda_C** and **Celda_G** models to perform co-clustering of genes into transcriptional modules and cells into subpopulations. A co-clustering topic model was previously developed called "Latent Dirichlet Co-Clustering" [5], in which each document is modeled as a mixture of document topics, each topic is a distribution over some paragraphs of the text, each of paragraph in the document is a mixture of word topics, and each word topic is a distribution over words. Here, we model each sample as a mixture of cellular subpopulations, each subpopulation as a mixture of transcriptional modules, and each transcriptional module as a mixture of genes. Note that we include the "hard-clustering" approach of **Celda_G** where each gene can only belong to a single transcriptional module.

#### 3.2 Generative process

1. Draw $\eta \sim \text{Dir}_L(\gamma)$
2. For each gene $g \in \{1..G\}$, draw $y_g \sim \text{Mult}(\eta)$
3. For each transcriptional module distribution $l \in \{1..L\}$ :
   (a) Define $Y_l = [y_g = l]_{g=1}^{G}$
   (b) Draw $\psi_l \sim \text{Dir}(\delta Y_l)$
4. For each sample $i \in \{1..S\}$, draw $\theta_i \sim \text{Dir}_K(\alpha)$
5. For each cell population $k \in \{1..K\}$, draw $\varphi_k \sim \text{Dir}_L(\beta)$
6. For each cell $j \in \{1..M_i\}$ in sample $i$:
   (a) Draw $z_{i,j} \sim \text{Mult}(\theta_i)$
   (b) For the $t$-th transcript in cell $j$ in sample $i$, $t \in \{1..N_{i,j}\}$:
      i. Draw $w_{i,j,t} \sim \text{Mult}(\varphi_{z_{i,j}})$
      ii. Draw $x_{i,j,t} \sim \text{Mult}(\psi_{w_{i,j,t}})$

**Description of parameters:**
$S$ is the number of samples.
$M_i$ is the number of cells in sample $i$.
$K$ is the number of cellular subpopulations
$L$ is the number of transcriptional modules.
$G$ is the number of genes.
$y_g$ is the hidden transcriptional module for gene $g$
$z_{i,j}$ is the hidden cell population for cell $j$ in sample $i$
$N_{i,j}$ is the number of transcripts for cell $j$ in sample $i$.
$w_{i,j,t}$ is the hidden transcriptional module for transcript $x_{i,j,t}$
$x_{i,j,t}$ is the $t^{th}$ transcript for cell $j$ in sample $i$.

Similar to the previous models, we refer to $\boldsymbol{\theta}$ as the "**Sample Probability (SP)**" matrix as it defines the probability of each cell population in each sample, $\boldsymbol{\varphi}$ as the "**Population Probability (PP)**" matrix as it defines the probability of each transcriptional module in each cell population, and $\boldsymbol{\psi}$ as the "**Gene Probability (GP)**" matrix as it defines the probability of each gene within each transcriptional module. The complete likelihood function is below followed by the plate diagram (Figure 3):

$$P(\boldsymbol{\eta}, \boldsymbol{\psi}, \boldsymbol{\theta}, \boldsymbol{\varphi}, \boldsymbol{Y}, \boldsymbol{Z}, \boldsymbol{W}, \boldsymbol{X} | \alpha, \beta, \gamma, \delta) = \\ P(\eta|\gamma) \prod_{g=1}^{G} P(y_g|\eta) \prod_{l=1}^{L} P(\psi_l|\delta, \boldsymbol{Y}) \prod_{i=1}^{S} p(\theta_i|\alpha) \prod_{k=1}^{K} P(\varphi_k|\beta) \prod_{j=1}^{M_i} p(z_{i,j}|\theta_i) \prod_{t=1}^{N_{i,j}} P(w_{i,j,t}|\varphi_{z_{i,j}}) P(x_{i,j,t}|\psi_{w_{i,j,t}}). \tag{51}$$

Figure 3: Plate diagram for cell and gene clustering model. Bold circles indicate given prior parameters and shaded circles indicate observed data.

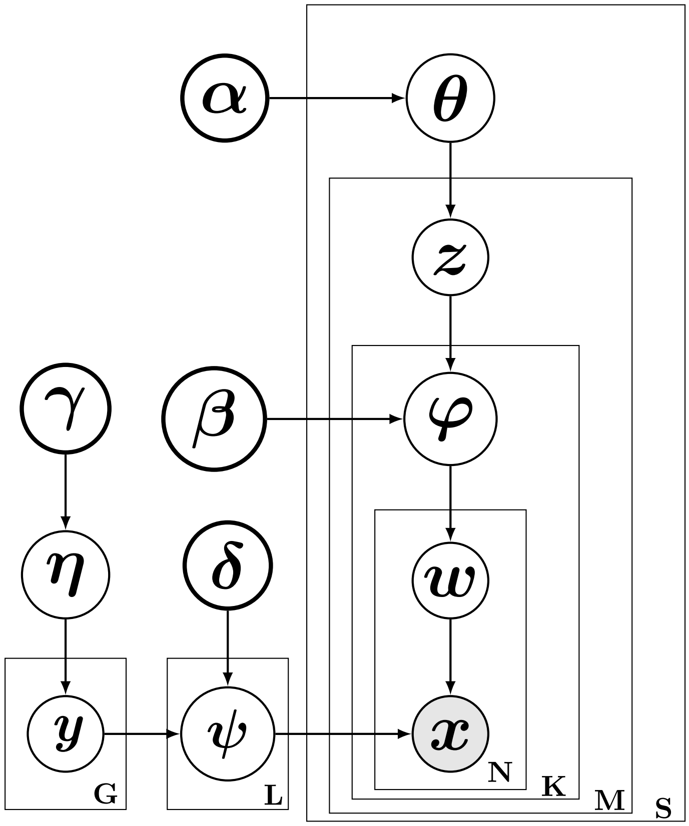


##### 3.3 Inference using collapsed Gibbs sampling

To build the collapsed Gibbs sampler, we next integrate out $\eta$, $\boldsymbol{\varphi}$, $\boldsymbol{\phi}$, $\boldsymbol{\psi}$, and $\boldsymbol{W}$:

$$\begin{aligned} P(\boldsymbol{Y},\boldsymbol{Z},\boldsymbol{X}|\alpha,\beta,\gamma,\delta) = &\int_{\eta}\int_{\boldsymbol{\psi}}\int_{\boldsymbol{\theta}}\int_{\boldsymbol{\varphi}} P(\eta|\gamma)\prod_{g=1}^{G}P(y_g|\eta)\prod_{l=1}^{L}P(\psi_l|\delta,Y_l)\prod_{i=1}^{S}p(\theta_i|\alpha)\prod_{k=1}^{K}P(\varphi_k|\beta)\prod_{j=1}^{M_i}p(z_{i,j}|\theta_i) \\ &\times\prod_{t=1}^{N_{i,j}}\left(\sum_{v=1}^{L}P(w_{i,j,t}=v|\varphi_{z_{i,j}})P(x_{i,j,t}=g|\psi_v)\right)d\boldsymbol{\varphi}d\boldsymbol{\theta}d\boldsymbol{\psi}d\eta. \end{aligned} \tag{52}$$

Using the procedure outlined in Celda_G, we can reduce the sum related to the marginalization of $\boldsymbol{W}$ by removing all components that are zero (i.e. all components where $w_{i,j,t} \neq y_g$):

$$P(\boldsymbol{Y},\boldsymbol{Z},\boldsymbol{X}|\alpha,\beta,\gamma,\delta)=\int_\eta\int_{\boldsymbol{\psi}}\int_{\boldsymbol{\theta}}\int_{\boldsymbol{\varphi}}P(\eta|\gamma)\prod_{g=1}^{G}P(y_g|\eta)\prod_{l=1}^{L}P(\psi_l|\delta,Y_l)\prod_{i=1}^{S}p(\theta_i|\alpha)\prod_{k=1}^{K}P(\varphi_k|\beta)\prod_{j=1}^{M_i}p(z_{i,j}|\theta_i) \times\prod_{t=1}^{N_{i,j}}P(w_{i,j,t}=y_g|\varphi_{z_{i,j}})P(x_{i,j,t}=g|\psi_{y_g})d\boldsymbol{\varphi}d\boldsymbol{\theta}d\boldsymbol{\psi}d\eta. \tag{53}$$

To integrate out the probabilities from the Dirichlet-multinomial distributions, we group terms related to $\eta$, $\boldsymbol{\theta}$, $\boldsymbol{\varphi}$, and $\boldsymbol{\psi}$:

$$\begin{aligned}P(\boldsymbol{Y},\boldsymbol{Z},\boldsymbol{X}|\alpha,\beta,\gamma,\delta)= \quad &\int_\eta P(\eta|\gamma)\prod_{g=1}^{G}P(y_g|\eta)d\eta\\ &\times\int_{\boldsymbol{\theta}}\prod_{i=1}^{S}p(\theta_i|\alpha)\prod_{j=1}^{M_i}p(z_{i,j}|\theta_i)d\boldsymbol{\theta}\\ &\times\int_{\boldsymbol{\varphi}}\prod_{k=1}^{K}P(\varphi_k|\beta)\prod_{i=1}^{S}\prod_{j=1}^{M_i}\prod_{t=1}^{N_{i,j}}P(w_{i,j,t}=y_g|\varphi_{z_{i,j}})d\boldsymbol{\varphi}\\ &\times\int_{\boldsymbol{\psi}}\prod_{l=1}^{L}P(\psi_l|\delta,Y_l)\prod_{i=1}^{S}\prod_{j=1}^{M_i}\prod_{t=1}^{N_{i,j}}P(x_{i,j,t}=g|\psi_{y_g})d\boldsymbol{\psi}.\end{aligned} \tag{54}$$

The integration for $\boldsymbol{\theta}$ and $\eta$ and follow the same procedures described in equations (13) and (34), respectively. Next, we can rearrange the components related to $\boldsymbol{\varphi}$ to focus on a single $\varphi_k$ as they are independent of one another. If we define $C_k$ to be the set of cells assigned to subpopulation $k$ and $C_k^i$ to be the set of cells assigned to population $k$ in sample $i$, then we have:

$$\int_{\boldsymbol{\varphi}}\prod_{k=1}^{K}P(\varphi_k|\beta)\prod_{i=1}^{S}\prod_{j=1}^{M_i}\prod_{t=1}^{N_{i,j}}P(w_{i,j,t}=y_g|\varphi_{z_{i,j}})d\boldsymbol{\varphi}=\prod_{k=1}^{K}\int_{\varphi_k}P(\varphi_k|\beta)\prod_{i=1}^{S}\prod_{j\in C_k^i}\prod_{t=1}^{N_{i,j}}P(w_{i,j,t}=y_g|\varphi_k)d\varphi_k. \tag{55}$$

Note that the index for $\varphi$ has been changed from $z_{i,j}$ to $k$ in $P(w_{i,j,t}=y_g|\varphi_k)$ because we have grouped all cells that have the same $k$ rather than ordering them by their index in sample $i$ (i.e. $\prod_{j=1}^{M_i}$). The series of multinomial probabilties for $\varphi_k$ can be re-written as:

$$\prod_{i=1}^{S}\prod_{j\in C_k^i}\prod_{t=1}^{N_{i,j}}P(w_{i,j,t}=y_g|\varphi_k)=\prod_{l=1}^{L}\varphi_{k,l}^{n_{(\cdot),(k),(V_l)}}, \tag{56}$$

where $n_{(\cdot),(k),(V_l)}$ represents the sum of counts across all samples for cells assigned to subpopulation $k$ for genes belonging to transcriptional module $l$. We can then integrate out $\varphi_k$ as follows according to the process described in equation (5):

$$\begin{aligned}\int_{\varphi_k}P(\varphi_k|\beta)\prod_{i=1}^{S}\prod_{j\in C_k^i}\prod_{t=1}^{N_{i,j}}P(w_{i,j,t}=y_g|\varphi_k)d\varphi_k&=\int_{\varphi_k}\frac{\Gamma(L\beta)}{\Gamma(\beta)^L}\prod_{l=1}^{L}\varphi_{k,l}^{\beta-1}\prod_{l=1}^{L}\varphi_{k,l}^{n_{(\cdot),(k),(V_l)}}d\varphi_k\\ &=\frac{\Gamma(L\beta)}{\Gamma(\beta)^L}\frac{\prod_{l=1}^{L}\Gamma(n_{(\cdot),(k),(V_l)}+\beta)}{\Gamma(\sum_{l=1}^{L}(n_{(\cdot),(k),(V_l)}+\beta))}.\end{aligned} \tag{57}$$

Next, we focus on the integration of $\boldsymbol{\psi}$. Let, $n_{(\cdot),(\cdot),g}$ represent the sum of counts for gene $g$ across all cells from all samples. The multinomial probabilities related to $\boldsymbol{\psi}$ can be re-written as:

$$\prod_{i=1}^{S}\prod_{j=1}^{M_i}\prod_{t=1}^{N_{i,j}} P(x_{i,j,t} = g|\psi_{y_g}) = \prod_{l=1}^{L}\prod_{v\in V_l} \psi_{l,v}^{n_{(\cdot),(\cdot),v}}, \tag{58}$$

where $\prod_{v\in V_l}$ represents a product over the set of genes assigned to transcriptional module $l$, similar to equation (37). Terms related to a specific $\psi_l$ can then be grouped and integreted over:

$$\begin{aligned}\int_{\boldsymbol{\psi}} \prod_{l=1}^{L} P(\psi_l|\delta, Y_l) \prod_{i=1}^{S}\prod_{j=1}^{M}\prod_{t=1}^{N_j} P(x_{i,j,t} = g|\psi_{y_g}) d\boldsymbol{\psi} &= \int_{\boldsymbol{\psi}} \prod_{l=1}^{L} \left( \frac{\Gamma(|V_l|\delta)}{\Gamma(\delta)^{|V_l|}} \prod_{v\in V_l} \psi_{l,v}^{\delta-1} \right) \prod_{l=1}^{L}\prod_{v\in V_l} \psi_{l,v}^{n_{(\cdot),(\cdot),v}} d\boldsymbol{\psi} \\ &= \prod_{l=1}^{L} \int_{\psi_l} \frac{\Gamma(|V_l|\delta)}{\Gamma(\delta)^{|V_l|}} \prod_{v\in V_l} \psi_{l,v}^{\delta-1} \prod_{v\in V_l} \psi_{l,v}^{n_{(\cdot),(\cdot),v}} d\psi_l \\ &= \prod_{l=1}^{L} \frac{\Gamma(|V_l|\delta)}{\Gamma(\delta)^{|V_l|}} \frac{\prod_{v\in V_l} \Gamma(n_{(\cdot),(\cdot),v} + \delta)}{\Gamma(\sum_{v\in V_l}(n_{(\cdot),(\cdot),v} + \delta))}.\end{aligned} \tag{59}$$

For clarity, the complete collapsed likelihood is as follows:

$$\begin{aligned}P(\boldsymbol{Y}, \boldsymbol{Z}, \boldsymbol{X}|\alpha, \beta, \gamma, \delta) = \quad & \frac{\Gamma(L\gamma)}{\Gamma(\gamma)^L} \times \frac{\prod_{l=1}^{L} \Gamma(|V_l| + \gamma)}{\Gamma(\sum_{l=1}^{L}(|V_l| + \gamma))} \\ & \times \prod_{i=1}^{S} \frac{\Gamma(K\alpha)}{\Gamma(\alpha)^K} \frac{\prod_{k=1}^{K} \Gamma(m_{i,k} + \alpha)}{\Gamma(\sum_{k=1}^{K}(m_{i,k} + \alpha))} \\ & \times \prod_{k=1}^{K} \frac{\Gamma(L\beta)}{\Gamma(\beta)^L} \frac{\prod_{l=1}^{L} \Gamma(n_{(\cdot),(k),(V_l)} + \beta)}{\Gamma(\sum_{l=1}^{L}(n_{(\cdot),(k),(V_l)} + \beta))} \\ & \times \prod_{l=1}^{L} \frac{\Gamma(|V_l|\delta)}{\Gamma(\delta)^{|V_l|}} \frac{\prod_{v\in V_l} \Gamma(n_{(\cdot),(\cdot),v} + \delta)}{\Gamma(\sum_{v\in V_l}(n_{(\cdot),(\cdot),v} + \delta))}.\end{aligned} \tag{60}$$

To perform Gibbs sampling, we will estimate the conditional distributions for the $\boldsymbol{Z}$ and $\boldsymbol{Y}$ indicator variables separately. For $\boldsymbol{Z}$, we have:

$$\begin{aligned}p(z_{i,j} = k|\boldsymbol{Z}_{-(i,j)}, \boldsymbol{Y}, \boldsymbol{X}, \alpha, \beta, \delta, \gamma) &= \frac{P(z_{i,j} = k, \boldsymbol{Z}_{-(i,j)}, \boldsymbol{Y}, \boldsymbol{X}|\alpha, \beta, \delta, \gamma)}{P(\boldsymbol{Z}_{-(i,j)}, \boldsymbol{Y}, \boldsymbol{X}|\alpha, \beta, \delta, \gamma)} \\ &\propto P(z_{i,j} = k, \boldsymbol{Z}_{-(i,j)}, \boldsymbol{Y}, \boldsymbol{X}|\alpha, \beta, \delta, \gamma).\end{aligned} \tag{61}$$

The first and last lines on the right side of equation (60) are invariant with respect to the configuration of cell population hidden variables $\boldsymbol{Z}$ and can be dropped. The second component can be simplified in the same manner as the corresponding component in the Celda_C model in equation (18). Therefore the final conditional distribution can be summarized as:

$$P(z_{i,j} = k, \boldsymbol{Z}_{-(i,j)}, \boldsymbol{Y}, \boldsymbol{X}|\alpha, \beta, \delta, \gamma) \propto (m_{i,k-(i,j)} + \alpha) \prod_{k=1}^{K} \frac{\Gamma(L\beta)}{\Gamma(\beta)^L} \frac{\prod_{l=1}^{L} \Gamma(n_{(\cdot),(k),(V_l)} + \beta)}{\Gamma(\sum_{l=1}^{L}(n_{(\cdot),(k),(V_l)} + \beta))}, \tag{62}$$

where $m_{i,(k)-(i,j)}$ is is the number of cells assigned to subpopulation $k$ in sample $i$ excluding the cell $z_{i,j}$. Importantly, the form of equation (62) is similar to that of the final line in equation (18) in Celda_C. The only difference is that Celda_C is performing the calculation over all genes whereas Celda_CG is performing the calculation over transcriptional modules. This can be elucidated by the fact that Celda_C uses $n_{(\cdot),(k),g}$ which is the sum of counts across all samples for cells in population $k$ for gene $g$, whereas Celda_CG uses $n_{(\cdot),(k),(V_l)}$ which also sums together counts across all genes in a transcriptional module $V_l$ in addition to summing across all cells in subpopulation $C_k$. In other words, the cell clustering occurs in a similar fashion as Celda_C, but on the reduced dimensional matrix of transcriptional modules rather than on the full matrix of genes.

Next, we will derive the Gibbs sampling equation for updates to $\boldsymbol{Y}$:

$$\begin{aligned} P(y_g = l|\boldsymbol{Y}_{-(g)}, \boldsymbol{Z}, \boldsymbol{X}, \alpha, \beta, \delta, \gamma) &= \frac{P(y_g = l, \boldsymbol{Y}_{-(g)}, \boldsymbol{Z}, \boldsymbol{X}|\alpha, \beta, \delta, \gamma)}{P(\boldsymbol{Y}_{-(g)}, \boldsymbol{Z}, \boldsymbol{X}|\alpha, \beta, \delta, \gamma)} \\ &\propto P(y_g = l, \boldsymbol{Y}_{-(g)}, \boldsymbol{Z}, \boldsymbol{X}|\alpha, \beta, \delta, \gamma). \end{aligned} \tag{63}$$

Similar to Celda_G, several components of equation (60) are invariant with respect to the configuration of the transcriptional module hidden variables $\boldsymbol{Y}$ and can be dropped. This includes the quantify $\frac{\Gamma(L\gamma)}{\Gamma(\gamma)^L}$ on the first line, the complete second line, the components $\frac{\Gamma(L\beta)}{\Gamma(\beta)^L}$ and $\Gamma(\sum_{l=1}^{L} n_{(\cdot),(k),(V_l)} + \beta)$ on the third line, and finally the quantity $\prod_{l=1}^{L}\prod_{v\in V_l}\Gamma(n_{(\cdot),(\cdot),v} + \delta)$ on the numerator of the last line. The final equation is as follows:

$$\begin{aligned} P(y_g = l, \boldsymbol{Y}_{-(g)}, \boldsymbol{Z}, \boldsymbol{X}|\alpha, \beta, \delta, \gamma) \propto \quad & \frac{\prod_{l=1}^{L}\Gamma(|V_l| + \gamma)}{\Gamma(\sum_{l=1}^{L}(|V_l| + \gamma))} \\ & \times \prod_{k=1}^{K}\prod_{l=1}^{L}\Gamma(n_{(\cdot),(k),(V_l)} + \beta) \\ & \times \prod_{l=1}^{L}\frac{\Gamma(|V_l|\delta)}{\Gamma(\delta)^{|V_l|}}\frac{1}{\Gamma(\sum_{v\in V_l}(n_{(\cdot),(\cdot),v} + \delta))}. \end{aligned} \tag{64}$$

This form of the equation is equivalent to equation (44) in **Celda_G** except for the fact that individual cells have been replaced with cell populations. In essense, all cells from the same subpopulation are summed together and the inference of transcriptional modules is performed on this reduced matrix. While **Celda_C** and **Celda_G** could be run separately to cluster cells and genes, **Celda_CG** will be significantly faster than running each of the other two models and is essential for understanding how each cell population can be described as a different combination of transcriptional modules.

The second and third terms in the above equation can be simplified in a similar manner as in **Celda_G** to further speed up updating $y_g$. Given the current gene $g$ is in module $l$, the second term can be simplified as

$$
\begin{aligned}
\prod_{k=1}^{K}\prod_{l=1}^{L}\Gamma(n_{(\cdot),(k),(V_l)}+\beta) &= \prod_{k=1}^{K}\left[\prod_{l=1}^{L}\Gamma(n_{(\cdot),(k),(V_l)}+\beta)\right] \\
&= \prod_{k=1}^{K}\left[\Gamma(n_{(\cdot),(k),(V_l)}+\beta)\prod_{l`:l`\neq l}\Gamma(n_{(\cdot),(k),(V_{l'})}+\beta)\right] \\
&= \prod_{k=1}^{K}\left[\Gamma(n_{(\cdot),(k),(V_l)}+\beta)\prod_{l`:l`\neq l}\Gamma(n_{(\cdot),(k),(V_{l'}^{-(g)})}+\beta)\right] \\
&= \prod_{k=1}^{K}\left[\frac{\Gamma(n_{(\cdot),(k),(V_l)}+\beta)}{\Gamma(n_{(\cdot),(k),(V_l^{-(g)})}+\beta)}\prod_{l`=1}^{L}\Gamma(n_{(\cdot),(k),(V_{l'}^{-(g)})}+\beta)\right] \\
&\propto \prod_{k=1}^{K}\left[\frac{\Gamma(n_{(\cdot),(k),(V_l)}+\beta)}{\Gamma(n_{(\cdot),(k),(V_l^{-(g)})}+\beta)}\right],
\end{aligned}
\tag{65}
$$

and the third term can be simplified as

$$
\begin{aligned}
\prod_{l=1}^{L}\frac{\Gamma(|V_l|\delta)}{\Gamma(\delta)^{|V_l|}}\frac{1}{\Gamma(\sum_{v\in V_l}(n_{(\cdot),(\cdot),v}+\delta))} &= \prod_{l=1}^{L}\frac{\Gamma(|V_l|\delta)}{\Gamma(\delta)^{|V_l|}}\times\prod_{l=1}^{L}\frac{1}{\Gamma(\sum_{v\in V_l}(n_{(\cdot),(\cdot),v}+\delta))} \\
&= \left[\prod_{l=1}^{L}\frac{\Gamma(|V_l|\delta)}{\Gamma(\delta)^{|V_l|}}\right]\times\left[\frac{1}{\Gamma(\sum_{v\in V_l}(n_{(\cdot),(\cdot),v}+\delta))}\prod_{l`:l`\neq l}\frac{1}{\Gamma(\sum_{v\in V_{l`}^{-(g)}}(n_{(\cdot),(\cdot),v}+\delta))}\right] \\
&= \left[\prod_{l=1}^{L}\frac{\Gamma(|V_l|\delta)}{\Gamma(\delta)^{|V_l|}}\right]\times\left[\frac{\Gamma(\sum_{v\in V_l^{-(g)}}(n_{(\cdot),(\cdot),v}+\delta))}{\Gamma(\sum_{v\in V_l}(n_{(\cdot),(\cdot),v}+\delta))}\prod_{l`=1}^{L}\frac{1}{\Gamma(\sum_{v\in V_{l`}^{-(g)}}(n_{(\cdot),(\cdot),v}+\delta))}\right] \\
&\propto \left[\prod_{l=1}^{L}\frac{\Gamma(|V_l|\delta)}{\Gamma(\delta)^{|V_l|}}\right]\times\left[\frac{\Gamma(\sum_{v\in V_l^{-(g)}}(n_{(\cdot),(\cdot),v}+\delta))}{\Gamma(\sum_{v\in V_l}(n_{(\cdot),(\cdot),v}+\delta))}\right],
\end{aligned}
\tag{66}
$$

where $V_l^{-(g)}$ is the total number of genes in module $l$ leaving out gene $g$. Notice that when gene $g$ is currently not in module $l$, $V_{l`}^{-(g)}$ is the same as $V_{l`}$.

Hence the full Gibbs sampling equation is simplified as:

$$
\begin{aligned}
P(y_g=l,\boldsymbol{Y}_{-(g)},\boldsymbol{Z},\boldsymbol{X}|\alpha,\beta,\delta,\gamma) \propto\ & \frac{\prod_{l=1}^{L}\Gamma(|V_l|+\gamma)}{\Gamma(\sum_{l=1}^{L}(|V_l|+\gamma))} \\
&\times\prod_{k=1}^{K}\left[\frac{\Gamma(n_{(\cdot),(k),(V_l)}+\beta)}{\Gamma(n_{(\cdot),(k),(V_l^{-(g)})}+\beta)}\right] \\
&\times\left[\prod_{l=1}^{L}\frac{\Gamma(|V_l|\delta)}{\Gamma(\delta)^{|V_l|}}\right]\times\left[\frac{\Gamma(\sum_{v\in V_l^{-(g)}}(n_{(\cdot),(\cdot),v}+\delta))}{\Gamma(\sum_{v\in V_l}(n_{(\cdot),(\cdot),v}+\delta))}\right].
\end{aligned}
\tag{67}
$$

After completing Gibbs sample and identifying the $\boldsymbol{Z}$ and $\boldsymbol{Y}$ with the highest probability, point estimates for the Dirichlet distribution probabilities can derived as described in the next section.

##### 3.4 Inference using point estimates

With a given sample of $\boldsymbol{Y}$ and $\boldsymbol{Z}$, we can derive point estimates for the Dirichlet distributions. For $\boldsymbol{\theta}$, we have:

$$\hat{\theta}_{i,k} = \frac{m_{i,k} + \alpha}{M_i + K\alpha}, \tag{68}$$

where $m_{i,(k)}$ is the number of cells assigned to population $k$ in sample $i$ and $M_i$ is the total number of cells in sample $i$. For $\boldsymbol{\varphi}$, we have:

$$\hat{\varphi}_{k,l} = \frac{n_{(\cdot),(k),(V_l)} + \beta}{n_{(\cdot),(k),(\cdot)} + L\beta}, \tag{69}$$

where $n_{(\cdot),(k),(V_l)}$ is the sum of counts across all samples across all cells belonging to subpopulaiton $k$ across all genes belonging to transcriptional module $l$ . $n_{(\cdot),(k),(\cdot)}$ is the sum of counts across all samples and across all cells belonging to subpopulation $k$ across all genes. For $\boldsymbol{\psi}$, we have:

$$\hat{\psi}_{l,g} = \frac{n_{(\cdot),(\cdot),g} + \delta}{n_{(\cdot),(\cdot),(V_l)} + |V_l|\delta}, \tag{70}$$

if $y_g = l$ (i.e. if gene $g$ is assigned to transcriptional module $l$). If $y_g \neq l$, then the posterior probability is 0. $n_{(\cdot),(\cdot),g}$ is the sum of counts across all samples and cells for gene $g$ and $n_{(\cdot),(\cdot),(V_l)}$ is the sum of counts all samples and cells across all genes belonging to module $l$.

Given $\hat{\boldsymbol{\theta}}$ and $\hat{\boldsymbol{\varphi}}$, we can identify the most likely cluster label for each $z_{i,j}$:

$$\begin{aligned} \hat{z}_{i,j} &= \text{argmax}_k \left\{ p(z_{i,j} = k | \boldsymbol{X}_{i,j}, \hat{\boldsymbol{\theta}}, \hat{\boldsymbol{\varphi}}) \right\} \\ &= \text{argmax}_k \left\{ \hat{\theta}_{i,k} \prod_{l=1}^{L} \hat{\varphi}_{k,l}^{n_{i,j,(V_l)}} \right\}, \end{aligned} \tag{71}$$

where $\boldsymbol{X}_{i,j}$ is the counts for cell $j$ in sample $i$ and $n_{i,j,(V_l)}$ is the sum of counts belonging to transcriptional module $l$ in cell $j$ in sample $i$. Note the form of this equation is similar to the point estimate for $Z$ in **Celda_C**, with the exception that genes have been collapsed into transcriptional modules. In this inference procedure, we alternate between estimating the Dirichlet distribution probabilities ($\hat{\boldsymbol{\theta}}$ and $\hat{\boldsymbol{\varphi}}$), the cell labels ($\hat{z}_{i,j}$), and the gene cluster labels $\boldsymbol{Y}$. $\boldsymbol{Y}$ is still estimated with Gibbs sampling conditioned on the point estimates for $\boldsymbol{Z}$ as decribed in the previous section. Since each cell is fully assigned to a subpopulation, this procedure is not truly expectation-maximization (EM). It more closely resembles the procedure used in K-means clustering, which is sometimes refered to as "hard" EM.

Although not explicitly specified in this model, we can also derive a "**Module Probability (MP)**" matrix containing the probability of each transcriptional module in each individual cell, which can be used in downstream analyses. This is performed by dividing the number of counts assigned to transcriptional module $l$ for cell $j$ in sample $i$ by the total number of counts for cell $j$:

$$MP_{i,j,l} = \frac{n_{i,j,(V_l)}}{N_{i,j}}. \tag{72}$$

##### 3.5 Perplexity

Perplexity was previously defined in equation (23). For the Celda_CG model, we define $\log(p(x))$ to be:

$$\log(p(x)) = \sum_{i=1}^{S} \sum_{j=1}^{M_j} \log \left[ \sum_{k=1}^{K} \theta_{i,k} \prod_{g=1}^{G} \left( \sum_{l=1}^{L} \varphi_{k,l} \psi_{l,g} \right)^{n_{i,j,g}} \right], \tag{73}$$

where $\theta_{i,k}$ is the probabiity of a cell belonging to a cell population $k$ in sample $i$, $\varphi_{k,g}$ is the probability of transcriptional module $l$ in cell population $k$, $\psi_{l,g}$ is the probability of gene $g$ in transcriptional module $l$, $n_{i,j,g}$ is the count of gene $g$ in cell $j$ in sample $i$, and $\eta_l$ is the probability of a gene being assigned to transcriptional module $l$. Note that the only non-zero $\psi_{l,g}$ will be where $y_g = l$ for gene $g$ and thus the sum of the equation can be simplified to:

$$\log(p(x)) = \sum_{i=1}^{S} \sum_{j=1}^{M_j} \log \left[ \sum_{k=1}^{K} \theta_{i,k} \prod_{g=1}^{G} \left( \varphi_{k,y_g} \psi_{y_g,g} \right)^{n_{i,j,g}} \right]. \tag{74}$$
